# ESMDisPred: A Structure-Aware CNN-Transformer Architecture for Intrinsically Disordered Protein Prediction

**DOI:** 10.64898/2026.01.22.701204

**Authors:** Md Wasi Ul Kabir, Ayon Dey, Farzeen Nafees, Md Tamjidul Hoque

## Abstract

Intrinsically disordered proteins (IDPs) lack stable three-dimensional structures, yet play vital roles in key biological processes, including signaling, transcription regulation, and molecular scaffolding. Their structural flexibility presents significant challenges for experimental characterization and contributes to diseases such as cancer and neurodegenerative disorders. Accurate computational prediction of IDPs is important for advancing research and drug discovery, structural biology, and protein engineering. In this study, we introduce ESMDisPred, a novel structure-aware disorder predictor that builds on the representational power of Evolutionary Scale Modeling-2 (ESM2) protein language models. ESMDisPred integrates sequence embeddings with structural information from the Protein Data Bank (PDB) to deliver state-of-the-art prediction accuracy. Model performance is further enhanced through feature engineering strategies, including terminal residue encoding, statistical summarization, and sliding-window analysis. To capture both local sequence motifs and long-range dependencies, we designed a hybrid CNN-Transformer architecture that balances convolutional efficiency with the representational power of self-attention. On CAID3 benchmarks, our latest model achieves ROC-AUC 0.895, AP 0.778, and a max F1 of 0.759, outperforming recent methods. Our results highlight the importance of integrating protein language model embeddings with explicit structural information for improved disorder prediction.

## 1.1 Introduction

Intrinsically disordered regions (IDRs) play essential roles in various biological processes but remain difficult to characterize experimentally due to their lack of stable tertiary structure. This has led to a growing reliance on computational methods for IDR prediction [1, 2]. Recent advances in deep learning and protein language models (PLMs) [3, 4] have significantly improved the accuracy and scalability of such predictions. These models learn representations from protein sequences, capturing both evolutionary and structural information without requiring handcrafted features. Among the current state-of-the-art methods, DisPredict3.0 [5] has demonstrated strong performance across multiple benchmarks. However, assessments from the CAID (Critical Assessment of protein Intrinsic Disorder) community challenge [6] reveal limitations in predictive performance, particularly in capturing subtle or transient disorder characteristics. This motivates the development of a more advanced and structure-aware prediction framework.

Several recent methods reflect similar efforts to push the boundaries of IDR prediction. For example, rawMSA-disorder, introduced by Mirabello and Wallner, utilizes raw multiple sequence alignments as input without relying on pre-computed features like PSSMs [7]. This architecture allows the network to learn disorder-relevant evolutionary patterns directly from the alignment data. DisoFLAG-IDR, developed by Pang and Liu, adopts a multi-task learning framework based on PLM embeddings and graph convolutional networks (GCNs) [8]. This model not only predicts disordered regions but also infers six functional classes of IDRs, including those involved in protein, DNA, RNA, lipid, and ion binding, as well as linker segments. The flDPnn3a model, part of the flDPnn series developed by the Kurgan Lab, combines sequence-derived features and global protein attributes. It employs an ensemble learning strategy to predict both general disorder and four functional IDR types, including MoRFs and linkers [9].

Beyond the models discussed above, several complementary approaches shape today’s IDR-prediction landscape. SPOT-Disorder2 and the flDPnn family remain strong deep-learning baselines; their ensemble designs still compete on CAID despite the shift to single-sequence PLMs [9, 10]. Lightweight single-sequence tools—SETH, LMDisorder, and Metapredict-v3—enable rapid proteome-scale screening with accuracy that rivals those of heavier profile-based pipelines [11]. Classical physics-inspired and consensus methods—IUPred3, AIUPred, and MobiDB-lite are known for their interpretability and long-IDR specificity [12–14]. Structure-proxy baselines derived from AlphaFold predictions provide computationally efficient disorder estimates that rank mid-table on CAID benchmarks. However, they generally lag behind purpose-trained PLM-based disorder predictors, as confidence scores were not explicitly optimized for disorder detection [15]. New methods such as PUNCH2 and task-focused binders (e.g., IPA variants, DeepDISObind) show gains for specialized aims like disordered binding-site localization [16–18].

Recent trends also favor lightweight models that maintain high accuracy while offering faster runtime. DisorderUnetLM, introduced by Kotowski et al., employs a U-Net architecture combined with ProtT5 embeddings. It demonstrated that single-sequence models can match or exceed profile-based methods in some contexts [19]. Another new approach is UdonPred-combined, which integrates continuous disorder signals from NMR-derived TriZOD data with binary disorder labels from DisProt [20]. Finally, EBIND-protein focuses on predicting disordered binding regions, specifically in MoRFs. By using PLM embeddings, EBIND-protein identifies binding-prone segments within IDRs [21].

In our earlier work, we introduced DisPredict3.0, the most recent iteration of the DisPredict series, which integrates evolutionary representations derived from protein language models to improve the prediction of intrinsically disordered regions (IDRs) [5]. This approach achieved the top ranking on the Disorder NOX dataset in CAID2. Building on this foundation, we now present ESMDisPred, a structure-aware disordered protein predictor that incorporates embeddings from the Evolutionary Scale Modeling-2 (ESM2) language model [3]. ESM2 is considered the SOTA language model and has demonstrated exemplary performance in protein structure prediction (ESMFold). To further improve predictive accuracy, we integrate feature-engineering strategies— including terminal residue encoding, statistical summarization, and sliding-window analysis. To capture both local sequence motifs and long-range dependencies, ESMDisPred employs a hybrid CNN-Transformer architecture that combines the efficiency of convolutional filters with the representational power of self-attention.

Comprehensive benchmarking on CAID3 datasets [22] demonstrates that ESMDisPred consistently outperforms existing state-of-the-art methods across multiple evaluation metrics, including AUC, F1 max, and average precision. These results demonstrate the importance of incorporating language-model-derived embeddings and structural awareness from the PDB into disorder prediction.

The remainder of this paper is organized as follows. We first describe the dataset curation process and present exploratory data analysis. This analysis reveals key disorder characteristics: the predominance of disordered residues in the N- and C-terminal regions, sequence-length distributions, and the relative proportions of ordered versus disordered states. We then discuss the machine learning methods explored in this study. We define the evaluation metrics used for performance assessment. Next, we describe protein representation using ESM2 embeddings. These findings directly inform our feature engineering strategies, including terminal-residue encoding, sequence-length filtering, and windowing techniques. We then detail the ESMDisPred architecture. Finally, we present comprehensive benchmarking results against state-of-the-art disorder predictors on CAID datasets.

### 1.2 Materials and Methods

#### 1.2.1 Dataset

We carefully curated the dataset from the DisProt database (prior to the 2023_12 release) [1, 2] to ensure no overlap between the training and testing sets. The initial dataset was composed of 2,845 proteins from the latest DisProt release. To improve the quality of the dataset, we removed long protein sequences that could introduce noise or bias into the analysis. In order to further refine the dataset and avoid redundancy, we applied a 25% sequence identity cutoff, ensuring that similar protein sequences were excluded [23]. This process resulted in a final training set consisting of 2,020 proteins, which together account for a total of 1,043,829 amino acids.

We analyze the distribution of amino acids (Figure 1 (b)) in the training dataset across four structural states: Order, Disorder, Transition State, and their combination (Disorder + Transition State). DisProt transition state annotation marks regions in a protein where disorder-to-order (or order-to-disorder) structural changes occur upon binding or environmental change [1, 24]. The majority of residues, approximately 81.41%, are classified as ordered as expected, while disordered and transition residues make up 16.26% and 2.32%, respectively. For model training and evaluation, we consider the annotation of disordered and transition states (totaling 18.58% of the dataset) as the positive class for prediction. This grouping acknowledges the functional and structural continuum between fully disordered and transitional regions. This analysis shows that the dataset is highly imbalanced (see Figure 1 (b)).

**Figure 1.**
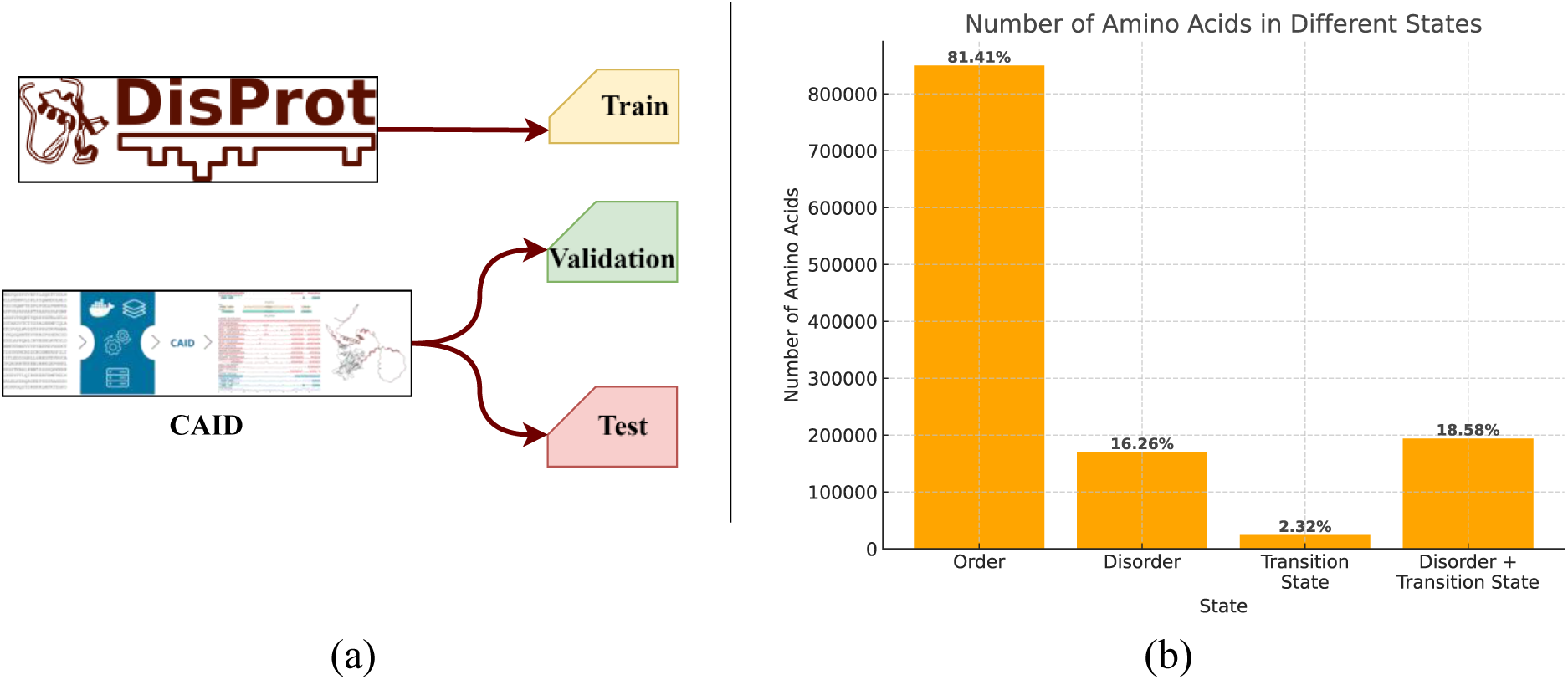
Overview of the dataset used for ESMDisPred model training and evaluation. (a) DisProt sequences are used for training, while CAID benchmark data are used for validation and testing. (b) Distribution of amino acids across different structural states (ordered, disordered, transition state) within the training dataset, showing strong class imbalance toward the ordered state.

For validation, we utilized two subsets from the CAID2 dataset: Disorder NOX (210 sequences) and Disorder PDB (348 sequences). The final model was tested using the CAID3 benchmark test datasets [25] to make sure there is no overlap between training, validation, and the test set.

#### 1.2.2 Exploratory Data Analysis

In this section, we explore the dataset to find relevant information that would help us increase the model’s accuracy. We begin by analyzing the distribution of protein sequences by length (Figure 2). Each bar in the figure represents the number of protein sequences corresponding to a particular sequence length, providing an overview of the length variation within the dataset. The analysis reveals that most protein lengths fall below 2000 amino acids. To reduce class imbalance, we exclude these longer sequences from the training dataset.

**Figure 2.**
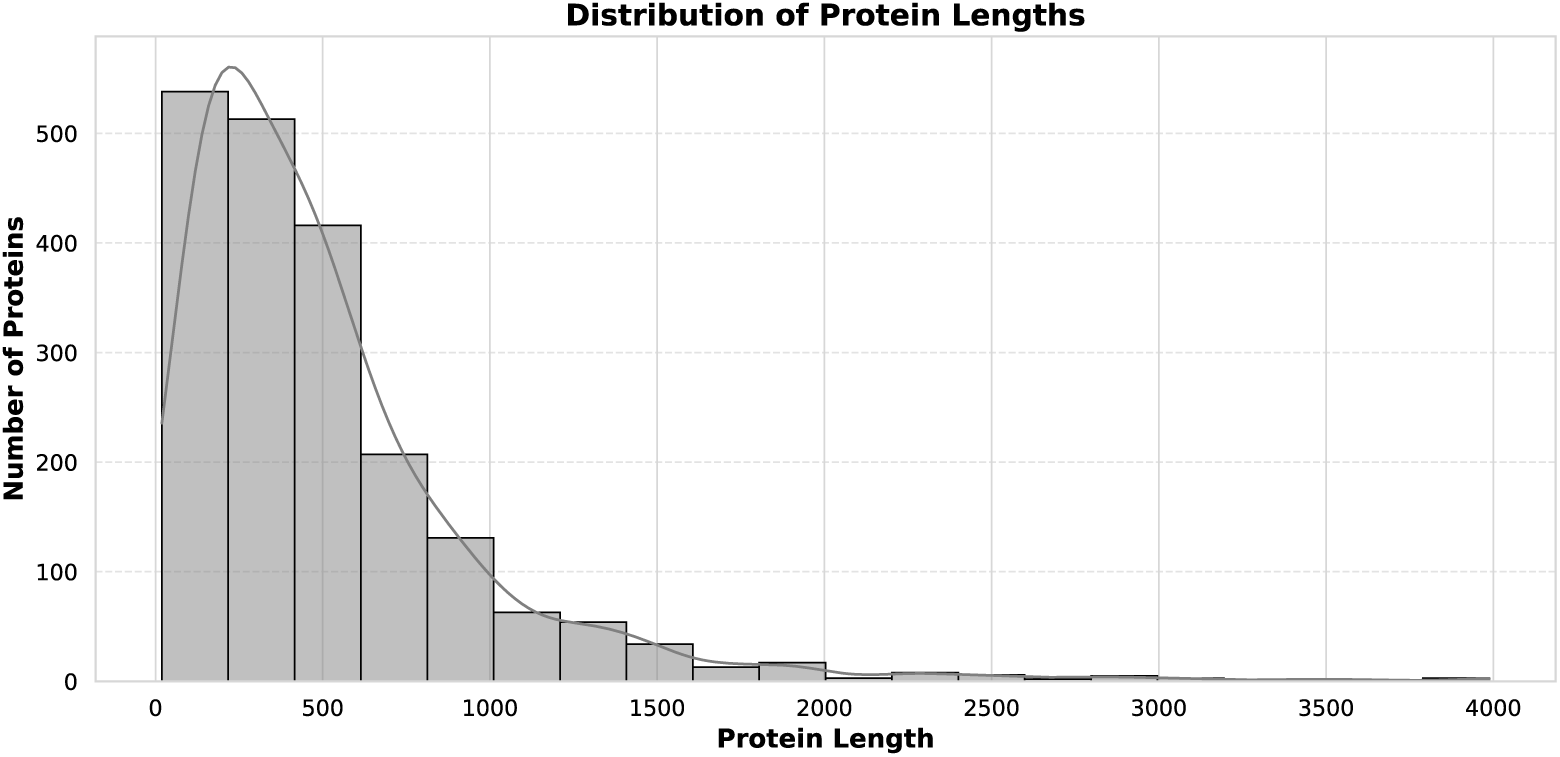
Distribution of protein sequences by length. The *x*-axis represents the length of the protein sequences (number of amino acids), and the *y*-axis shows the number of proteins corresponding to each sequence length. The majority of protein sequences in the dataset fall between 0 and 500 amino acids, with a sharp decline in the number of sequences as the length increases beyond 500 amino acids.

We also explore the distribution of disordered regions in proteins from the training set (Figure 3). Most disordered residues are concentrated in the N- and C-terminal regions. This pattern suggests that intrinsic disorder is not randomly distributed. Incorporating this context into predictive models by highlighting terminal regions can help improve disorder prediction.

**Figure 3.**
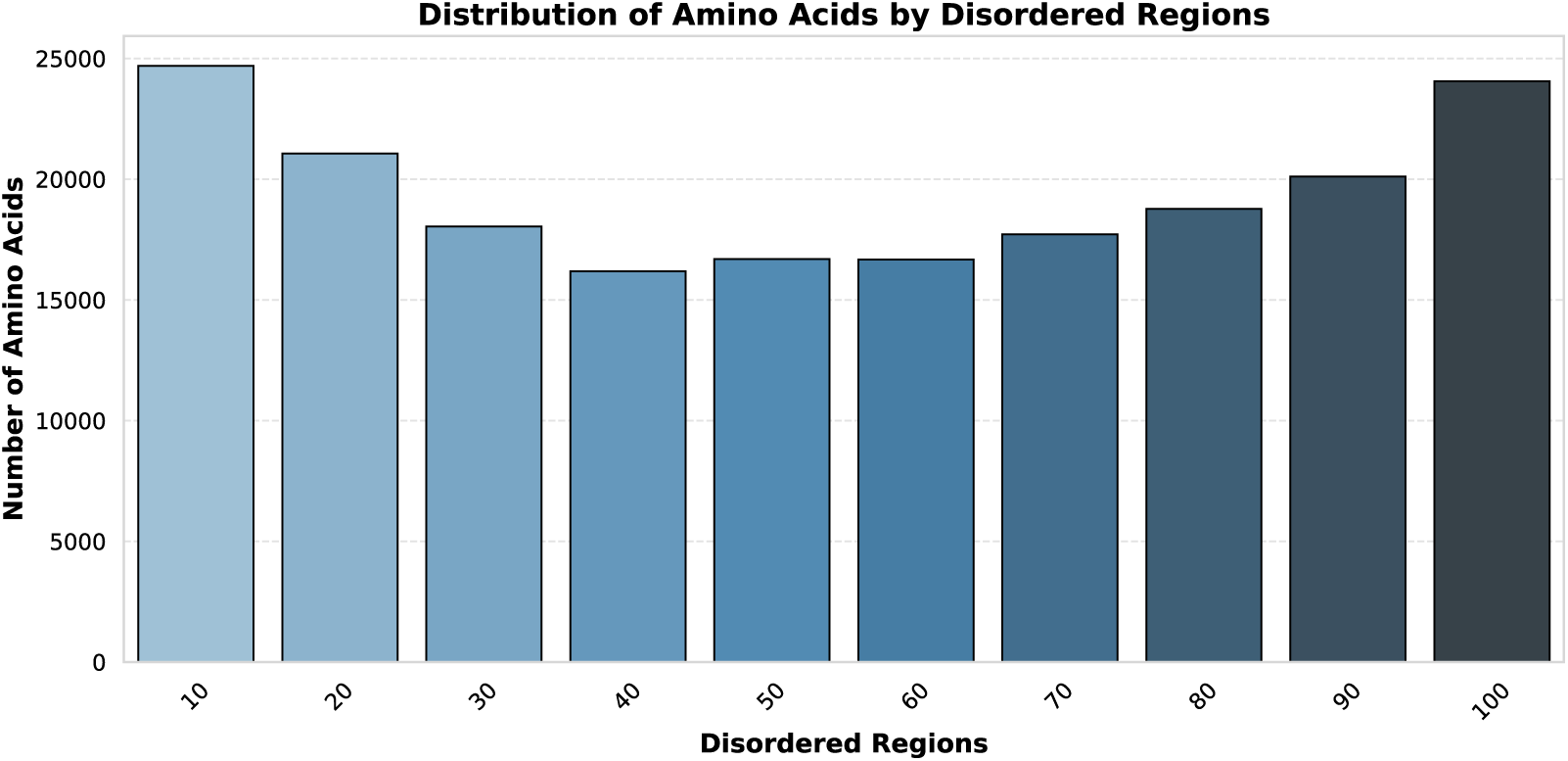
The distribution of disordered amino acids across protein sequence positions. The *x*-axis represents relative positions along the protein sequence, divided into 10 equal parts (from 10% to 100%). The *y*-axis indicates the total number of disordered amino acids observed in each segment across all proteins. The plot reveals that disordered amino acids are most frequently found at the N-terminal (10%) and C-terminal (100%) ends of proteins, while middle regions (30%–70%) contain fewer disordered residues.

Figure 4 shows the distribution of proteins by the percentage of disordered regions. As expected, the majority of proteins contain a low proportion of disordered residues, indicating that intrinsic disorder is often confined to specific regions rather than spread throughout the protein. This supports the notion that disorder commonly plays a localized role. However, the distribution also reveals a subset of proteins with a high percentage of disordered regions, including proteins that are almost entirely disordered. These fully disordered proteins likely perform specialized functions, such as signaling, molecular recognition, or forming dynamic complexes.

**Figure 4.**
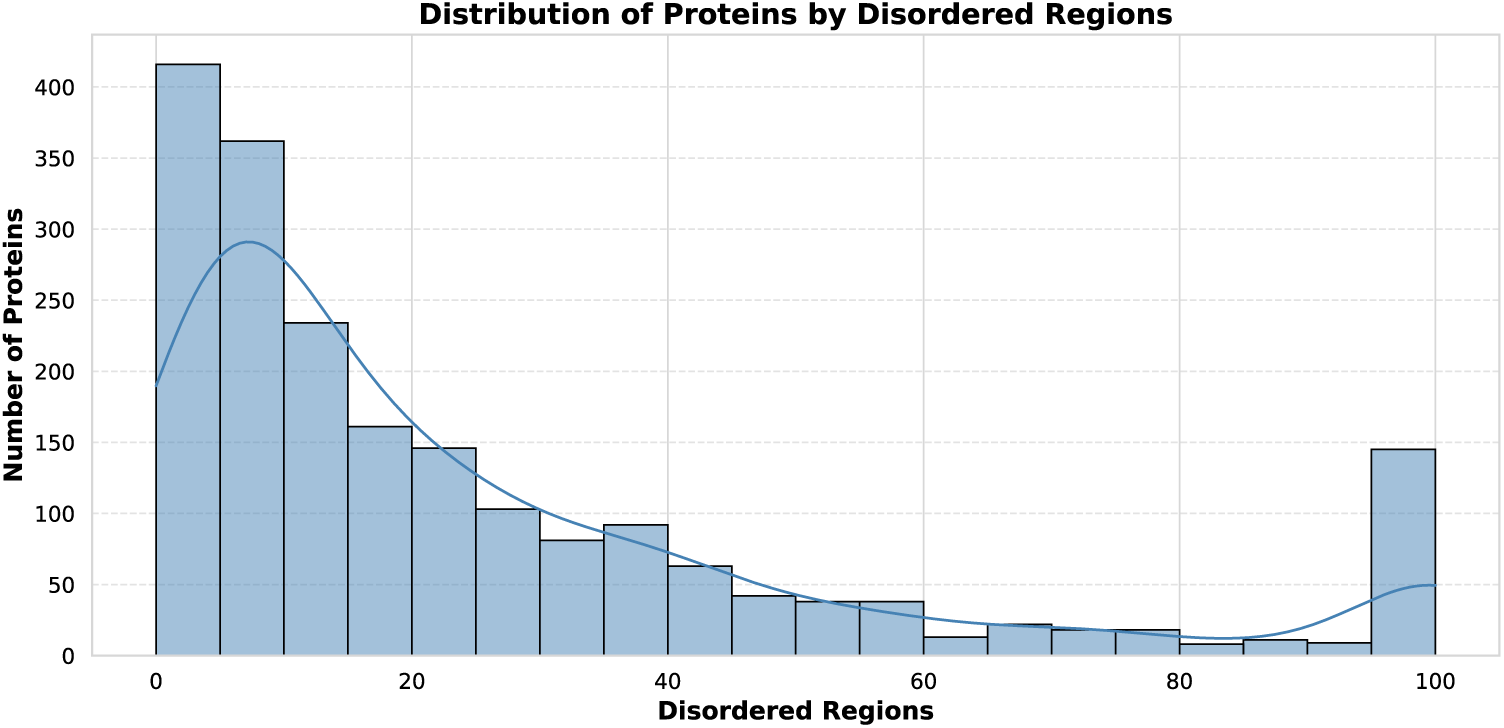
The distribution of proteins based on the percentage of disordered regions. The *x*-axis represents the percentage of disordered regions in proteins, while the *y*-axis shows the number of proteins in each percentage range. The majority of proteins have a low percentage of disordered regions (0-20%), with a sharp decline as the percentage of disorder increases. There is a noticeable peak at 100%, indicating that a substantial number of proteins are fully disordered.

The insights obtained from the exploratory analysis guide the next stages of model development. Specifically, identifying the sequence length distribution helps us address class imbalance by filtering out extreme outliers, while the observed enrichment of disorder at the N- and C-termini highlights the importance of positional information. Based on these findings, we performed targeted feature selection and engineering to capture sequence length variability, terminal region characteristics, and disorder proportions. We discuss these steps in detail in the Feature Selection and Engineering sections.

#### 1.2.3 Machine Learning methods

We explore a range of machine learning (ML) methods to have a baseline for the disorder predictor, including ensemble tree algorithms, and deep learning architectures. For instance-based learning, we used KNeighborsClassifier (KNN) [26]. This method classifies a sample based on the majority class of its nearest neighbors in feature space. We also studied several ensemble tree-based methods. RandomForestClassifier [27] constructs multiple decision trees using bootstrapped samples and randomly selected feature subsets. This bagging approach enhances generalization and reduces overfitting by reducing variance error from the average decision. ExtraTreesClassifier [28] builds an ensemble of completely randomized trees, introducing further randomness in the choice of split thresholds.

We evaluated boosting-based ensemble methods using four algorithms. AdaBoostClassifier [29], which combines weak learners in sequence, giving more weight to misclassified instances. XGBoost [30] improves classical gradient boosting through second-order optimization, regularization, and fast tree construction. LightGBM [31] incorporates Gradient-based One-Side Sampling (GOSS) and Exclusive Feature Bundling (EFB) to improve efficiency on large datasets. CatBoost [32] addresses categorical feature handling using ordered boosting and target statistics, reducing overfitting and bias.

In exploring deep learning techniques, we trained our dataset with the CategoryEmbeddingModel [33], a multilayer perceptron that uses learnable embeddings for categorical features. This approach replaces one-hot encodings with dense, continuous vectors that the model updates during training. We evaluated TabTransformer [34], which applies transformer layers to contextualize categorical embeddings before combining them with numerical features in a fully connected network. We also used TabNet [35], which introduces a sequential attention mechanism for selecting relevant features at each decision step. DANet [36] incorporates an Abstract Layer to construct higher-order feature interactions, aided by shortcut paths from input to deeper layers to preserve original feature representations.

We also evaluated GANDALF [37], a deep learning model based on Gated Recurrent Units (GRUs), adapted for tabular data. It introduces the Gated Feature Learning Unit (GFLU), which performs implicit feature selection through gating mechanisms.

In addition, we designed a CNN-Transformer hybrid architecture for per-residue disorder prediction (Algorithm 2). This model combines convolutional layers for capturing local motifs with Transformer encoders for long-range sequence dependencies. A full description of this model is provided in Section 1.4. The performance of all models on the validation set is reported in the Results section.

#### 1.2.4 Evaluation Metrics for Performance Assessment

Given the significant class imbalance in the disorder dataset, as illustrated in the Dataset section, we evaluate model performance using three key metrics: Area Under the Curve (AUC), Average Precision (AP), and the maximum F1 score (F1 max).

##### AUC (Area Under the ROC Curve)

AUC measures the model’s ability to rank positive instances higher than negative ones. It is computed as the area under the receiver operating characteristic (ROC) curve, which plots the true positive rate (TPR) against the false positive rate (FPR) (see Eq. (1)). The AUC value ranges from 0 to 1, where 1.0 indicates perfect discrimination, and 0.5 represents random performance.

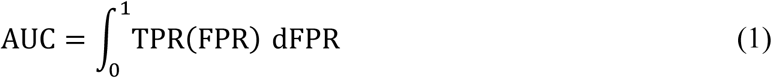

##### Average Precision Score (AP)

Average Precision summarizes the precision-recall curve and is defined as the weighted mean of precisions at each threshold (see Eq. (2)) where recall increases.

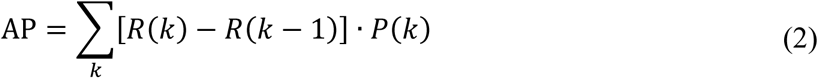

where, *P*(*k*) and *R*(*k*) denote the precision and recall at the *k* threshold. AP is informative for imbalanced datasets, where it reflects the model’s ability to retrieve relevant instances.

##### F1 max (Maximum F1 Score)

F1 score is the harmonic mean of precision and recall (see Eq. (3)):

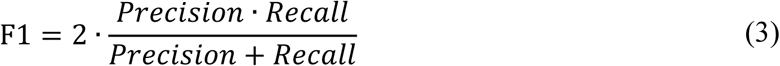

F1 max is the maximum F1 score obtained by varying the classification threshold *t* over a set of possible thresholds *T* (see Eq. (4)):

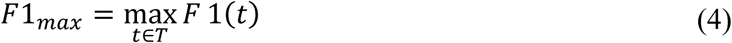

#### 1.2.5 Protein Representation

We represent proteins using protein language models (PLMs)—deep learning models inspired by natural language processing (NLP) techniques. These models treat amino acid sequences as a form of “biological language,” where sequences of residues carry implicit structural, functional, and evolutionary information. By training on large-scale protein databases such as UniProt, these models learn to capture sequence patterns, residue dependencies, and contextual representations [3, 38]. At the core of most PLMs lies the transformer architecture, originally developed for NLP tasks. Models like ESM (Evolutionary Scale Modeling), ProtBert, TAPE, and MSA Transformer have demonstrated the ability to learn high-quality embeddings of protein sequences [4, 39, 40]. These embeddings encode features such as secondary structure, disorder, binding affinity, and functional domains.

Protein language models are powerful because they are unsupervised, scalable, and general-purpose [22, 40]. Once trained, their learned embeddings can be fine-tuned or directly applied to a wide range of biological prediction tasks, often outperforming traditional methods [4, 39]. Their interpretability is still under exploration, but attention maps and latent representations offer insights into biologically meaningful patterns. As PLMs continue to scale and integrate structural supervision (as seen in ESMFold and AlphaFold-style models) [40–42], they are aiding in protein function annotation, design, and drug discovery [40, 41].

In this study, we extract embeddings from the ESM family of models, developed by Meta AI (formerly Facebook AI Research), which are large-scale Transformers trained on hundreds of millions of protein sequences using masked-language modeling. ESM embeddings have shown strong performance on tasks such as remote homology detection, structure prediction, and zero-shot classification. The most recent ESM2 and ESMFold models push this further by integrating language modeling with structure prediction, achieving near AlphaFold-level performance on certain benchmarks [40–42]. MSA Transformer (another variant) incorporates multiple sequence alignments as input, allowing the model to directly learn from evolutionary relationships across homologous sequences [39]. In addition to ESM2 language model embeddings, we incorporated disorder-specific features from our previously developed tool, DisPredict3.0 [5].

### 1.3 Feature Selection, Engineering, and Pre- and Post-processing

We integrated multiple types of features derived from state-of-the-art approaches, including protein language model embeddings, predicted structural representations, and sequence-based descriptors. To evaluate the relative importance of each feature type, we engineered individual features and measured the resulting changes in predictive performance. This ablation analysis quantified the specific contribution of each feature category to overall model accuracy.

The pipeline begins with data filtering to ensure quality. We removed proteins longer than 2000 amino acids and excluded proteins containing missing residues—amino acids present in the sequence but lacking atomic coordinates in PDB structures. These preprocessing steps reduce noise and class imbalance in the training data. The feature engineering strategy in ESMDisPred involves three key components: (1) disorder predictions generated by DisPredict3.0, (2) protein sequence embeddings extracted from ESM2, and (3) terminal residue annotations encoding each residue’s relative position within the sequence. These features are then transformed into statistical descriptors using techniques such as mean, variance, skewness, and kurtosis. Additionally, we apply sliding-window analysis to capture local sequence context. These transformations provide rich input representations that enhance model accuracy.

To improve residue-level prediction stability and reduce noise in the output probabilities, we evaluated three probability-smoothing methods: moving average, exponential moving average, and Gaussian filtering. These techniques refine raw predictions by incorporating information from neighboring residues, producing more coherent disorder profiles. All performance evaluations of preprocessing, feature engineering, and post-processing strategies are reported on the validation set to ensure consistent comparison. The following subsections discuss the impact of each technique.

#### 1.3.1 Effect of Large Protein Sequences

We first explore the impact of excluding large proteins on the performance of the model. As shown in Figure 5, filtering out large proteins based on sequence length thresholds affects AP, AUC, and F1 max metrics. The base model, which includes all proteins regardless of size, serves as a reference point. Removing proteins longer than 2000 residues yields the most performance gains, with AP improving to over 11% and F1 max reaching around 7%, resulting in the highest average performance improvement among all tested conditions. Excluding proteins above 2500 residues also boosts performance, though slightly less effectively. Conversely, applying a more aggressive filter by removing proteins longer than 1500 residues leads to weaker improvements. These findings indicate that eliminating extremely long proteins helps mitigate noise during model training and reduce imbalance in the dataset.

**Figure 5.**
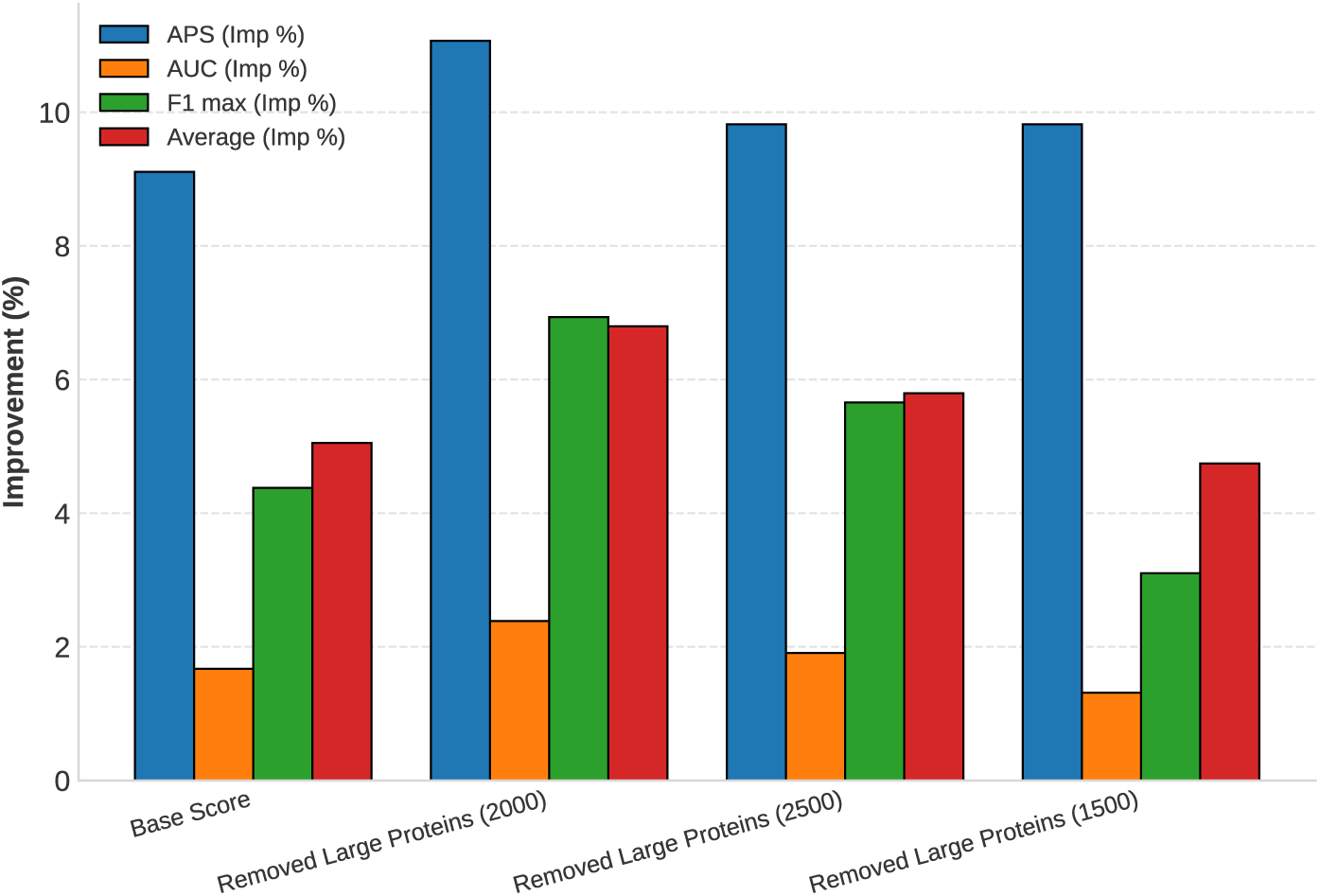
Impact of removing large proteins on various performance metrics. The percentage improvements in AP, AUC, F1 max, and the overall average performance are shown for different thresholds of protein length removal (1500, 2000, and 2500 amino acids) compared to the base model score. Excluding long proteins increases performance across all metrics, with the best overall gains achieved by removing sequences greater than 2000 amino acids.

#### 1.3.2 Effect of ESM2 Statistics

We further analyzed the ESM2-derived statistical features, including metrics such as sequence length, mean, standard deviation, variance, and higher-order moments (e.g., skewness and kurtosis) to assess their contribution to predictive performance. These statistics provided complementary information that improved the model’s robustness. Figure 6 illustrates the effect of incorporating ESM2-derived features on model performance. Incorporation of raw ESM2 embeddings leads to notable improvements, particularly in AP (∼12.2%) and F1 max (∼8%), demonstrating the utility of pretrained language model representations in capturing relevant protein sequence information. Further refinement using ESM2 statistical features results in the highest observed gains, with AP nearing 16% improvement and F1 max exceeding 11%. These results highlight that not only the inclusion of ESM2 features but also their statistical summarization can enhance the model’s predictive performance.

**Figure 6.**
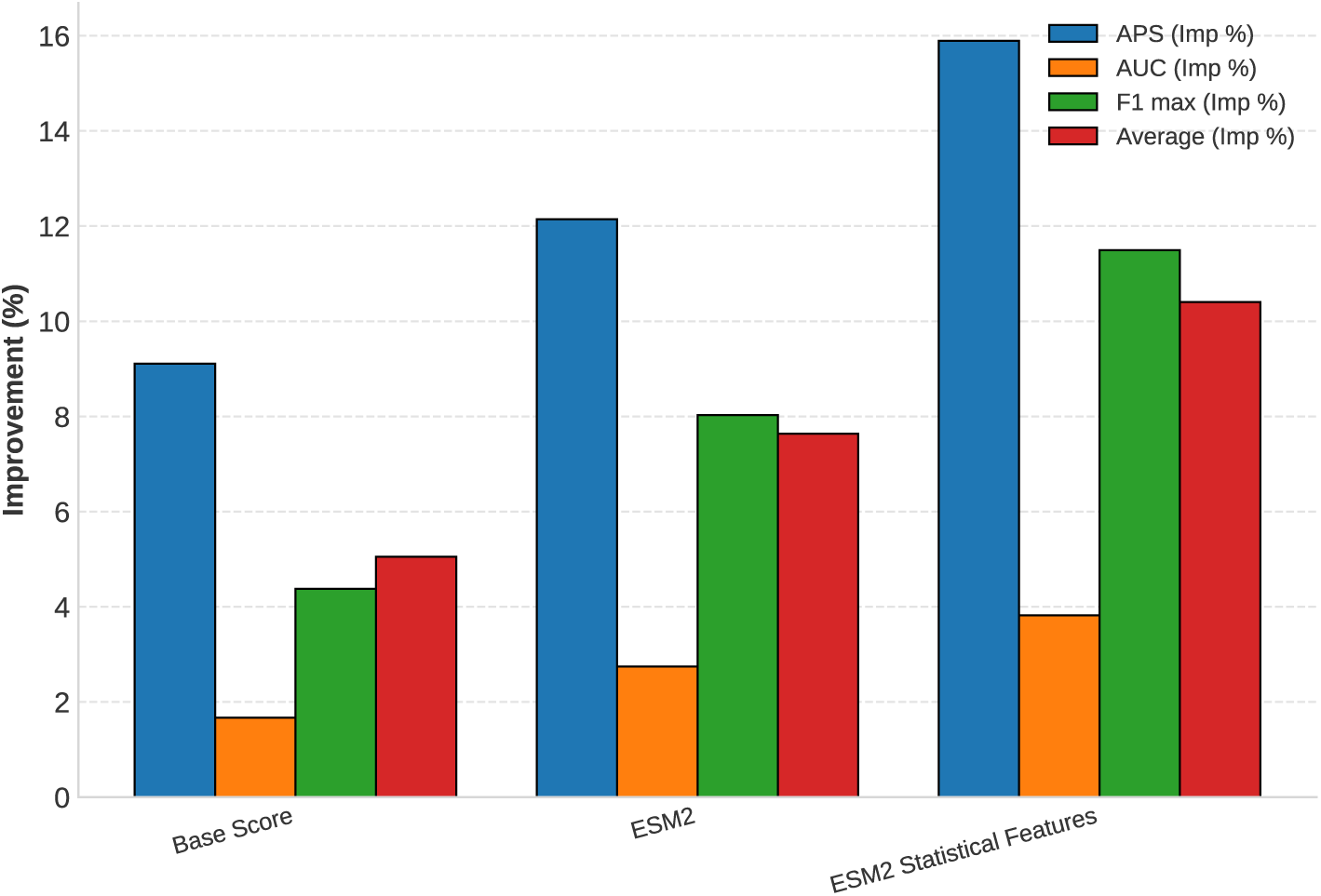
Performance improvements resulting from integrating ESM2 embeddings and ESM2-derived statistical features into the model.Adding ESM2 embeddings and ESM2-derived statistical features improves all metrics, and combining embeddings with statistical features yields the largest overall gains.

#### 1.3.3 Effect of Terminal Values

Exploratory data analysis revealed that disordered residues are predominantly concentrated in the N- and C-terminal regions of protein sequences. This observation motivated us to explicitly include terminal residue information in our feature set. Incorporating these regions led to improved predictive accuracy, particularly for proteins exhibiting high intrinsic disorder near sequence termini. Figure 7 illustrates two encoding schemes applied to a protein sequence: a positive-negative annotation from −1 to 1 (shown in blue) and a positive-symmetric annotation from 1 to 1 (shown in green). Each residue is assigned a numerical value based on its relative position, with central residues receiving values near zero and terminal residues approaching the extremes. Positive and negative encoding provide a continuous, direction-aware representation ideal for terminal residue annotation. In contrast, the positive symmetric encoding reflects distance from the sequence center without a directional sign.

**Figure 7.**
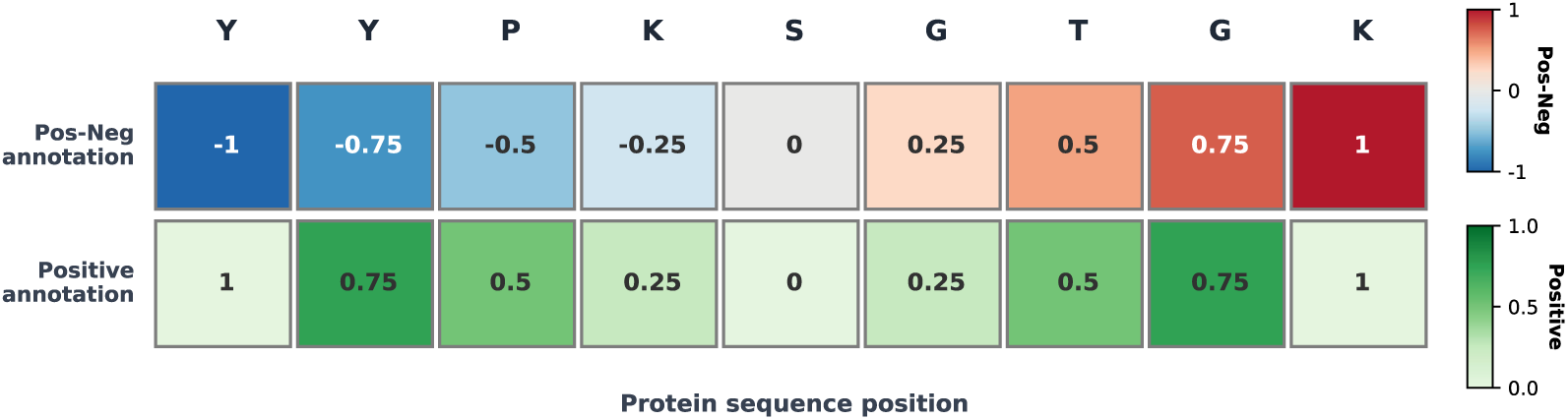
Example of length-normalized positional annotation for terminal-residue encoding (illustrated for a sequence of length L = 9). Each residue is assigned a value based on its relative position along the sequence using two schemes: (i) a signed encoding in [−1,1] (blue) that distinguishes the N-terminus (negative) from the C-terminus (positive) and varies linearly with position, and (ii) a symmetric encoding in [0,1] (green) that reflects proximity to either terminus, with values increasing toward the sequence ends.

Figure 8 examines the impact of terminal residue annotations on predictive performance. The base model, which lacks any positional encoding of terminal residues, shows the lowest performance. The “Terminal Pos-Neg” yields the highest improvement in AP (∼11.8%) and F1 max (∼8%). This indicates that emphasizing both termini enhances the model’s ability to capture biologically relevant positional cues. In contrast, the “Terminal Positive-Only” provides a modest improvement over the base but underperforms compared to the symmetric scheme.

**Figure 8.**
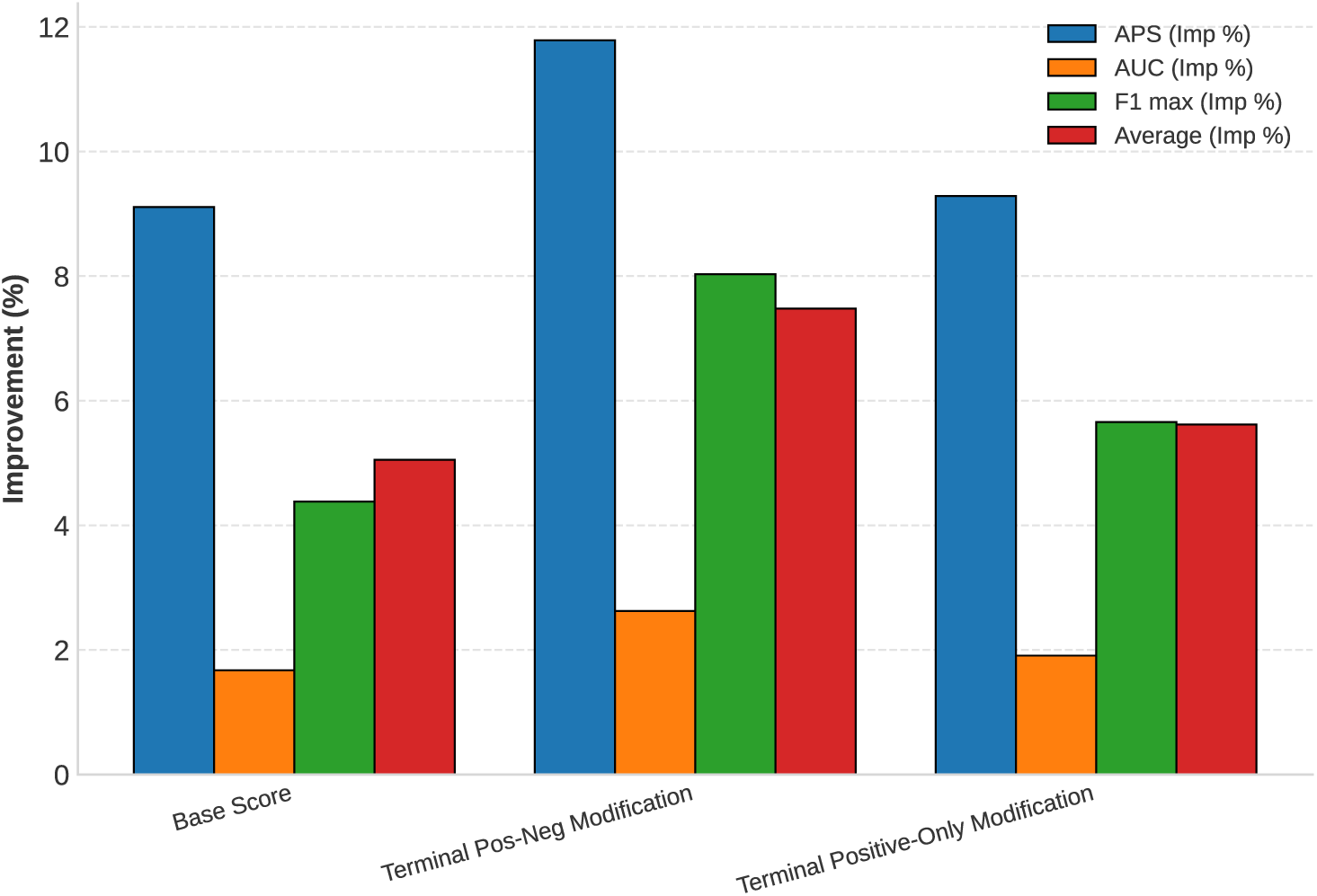
Effect of incorporating terminal residue annotations on model performance. The figure shows the percentage improvements relative to the baseline for two terminal-encoding schemes: a signed N-to-C annotation (Pos–Neg) and a symmetric positive-only annotation. Both schemes improve performance, with the signed Pos–Neg encoding yielding the overall better results.

#### 1.3.4 Effect of Removing Missing Residues

In our training dataset, we have residues (amino acids) present in the sequence but lacking atomic coordinates in experimentally determined structures (e.g., Protein Data Bank (PDB) entries). These missing residues often result from unresolved regions due to intrinsic flexibility, truncation, or experimental limitations. Although commonly treated as disordered, their status is ambiguous and can introduce noise into training data. To address this, we studied the effect of these residues on predictive performance. Figure 9 illustrates the effect of removing missing residues on performance metrics. Compared to the base model, removing proteins with missing residues led to a marked increase in predictive performance. Specifically, the Average Precision Score (AP) increased from ∼9% to over 12.5%, indicating improved ranking of disordered residues. The average of all metrics also improved, rising from ∼5% to ∼6%. Including missing residues can introduce inconsistencies between sequence and structure, degrade feature reliability, and mislabel flexible but ordered regions. These findings show the importance of data quality and improves the disorder prediction accuracy.

**Figure 9.**
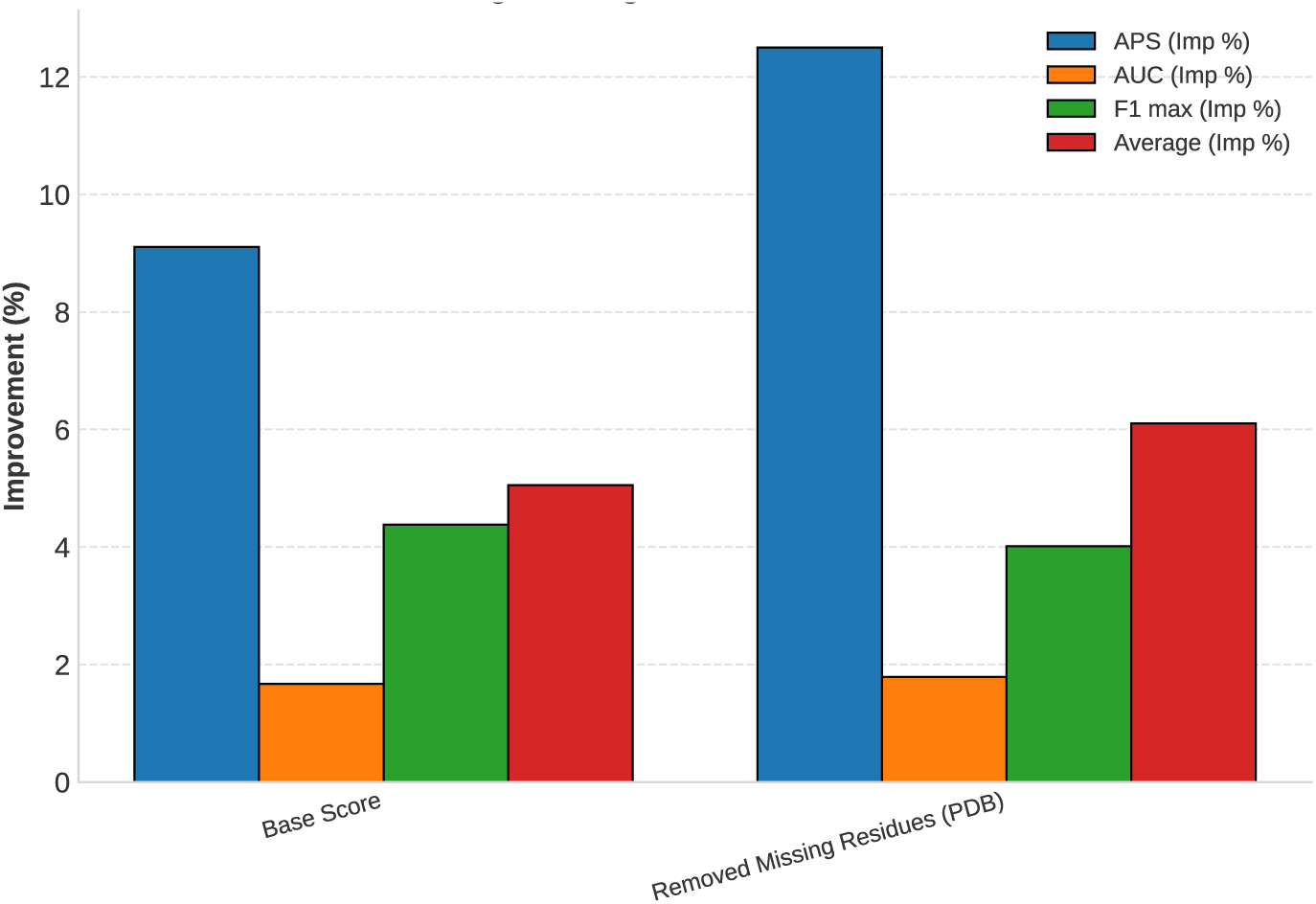
Effect of removing residues with missing structural coordinates (PDB) on model performance. Percentage improvements in APS, AUC, max F1, and the overall average are shown relative to the baseline after excluding amino-acid positions lacking resolved coordinates. Removing missing residues improves performance across all metrics.

#### 1.3.5 Effect of Windowing

The windowing technique is a widely used method that involves moving a fixed-length subsequence along the protein sequence to capture the local contextual information surrounding each residue [43–47]. Figure 10 illustrates the windowing technique on a protein sequence. In the left panel, the model processes the entire sequence as a whole, extracting features globally at each residue position. While this approach captures overall sequence information, it may miss important local patterns, particularly those associated with intrinsic disorder. The right panel shows the windowing strategy: a window of length 3 is moved along the sequence to form overlapping triplets (V–K–G, K–G–L,…). For each triplet, the residue-level feature vectors are concatenated to create a local representation (conceptually 3*d*). The dimensionalities shown in the figure (e.g., 96 and 288) are for illustration only; in our model, each residue is represented by a 1480-dimensional feature vector. This method enables the model to focus on local context and short-range dependencies, which are often critical for detecting regions of disorder. By aggregating the features from multiple overlapping windows, the model builds a localized representation of the sequence.

**Figure 10.**
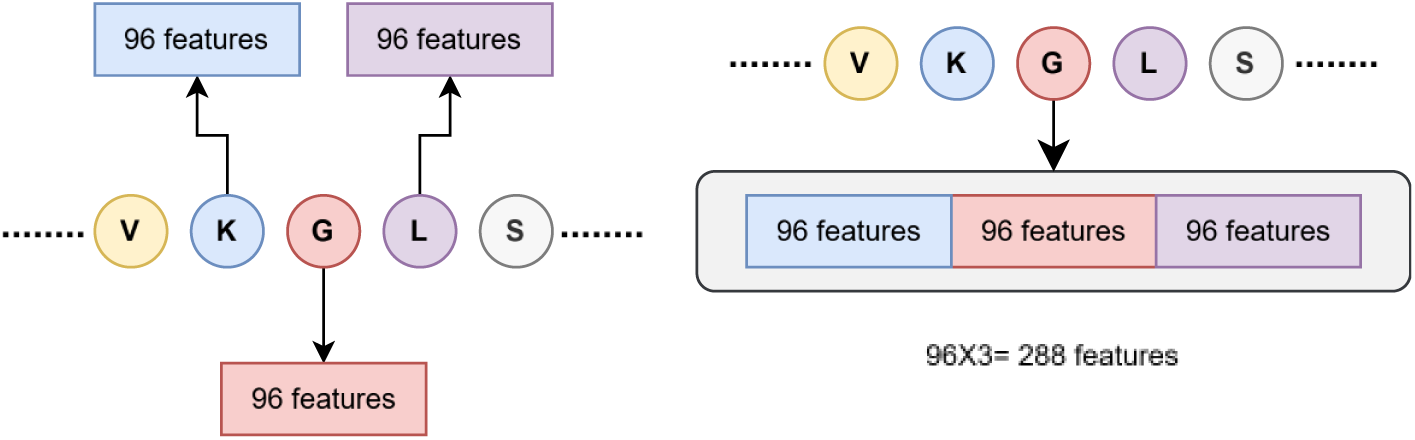
Illustration of window-based feature extraction from protein sequences. Each amino acid residue in a sequence (e.g., VKGLS) is represented by a set of statistical features (e.g., 96-dimensional feature vectors) derived from a sliding window approach. The window encompasses neighboring residues to capture local context, resulting in a feature representation that encodes local physicochemical and contextual information.

Applying windowing techniques to all features would increase dimensionality substantially and could lead to overfitting and degraded generalization due to the curse of dimensionality. To mitigate this, we applied the windowing technique selectively to features where local context is most informative, such as disorder probabilities, and ESM2-derived statistical features. Figure 11 presents the impact of applying windowing techniques with varying window sizes on the performance of selected features. The base score, which does not use windowing, shows the lowest performance across all metrics, whereas, the window size 7 yielded the highest overall improvement in AP (16%) and F1 max (11.5%).

**Figure 11.**
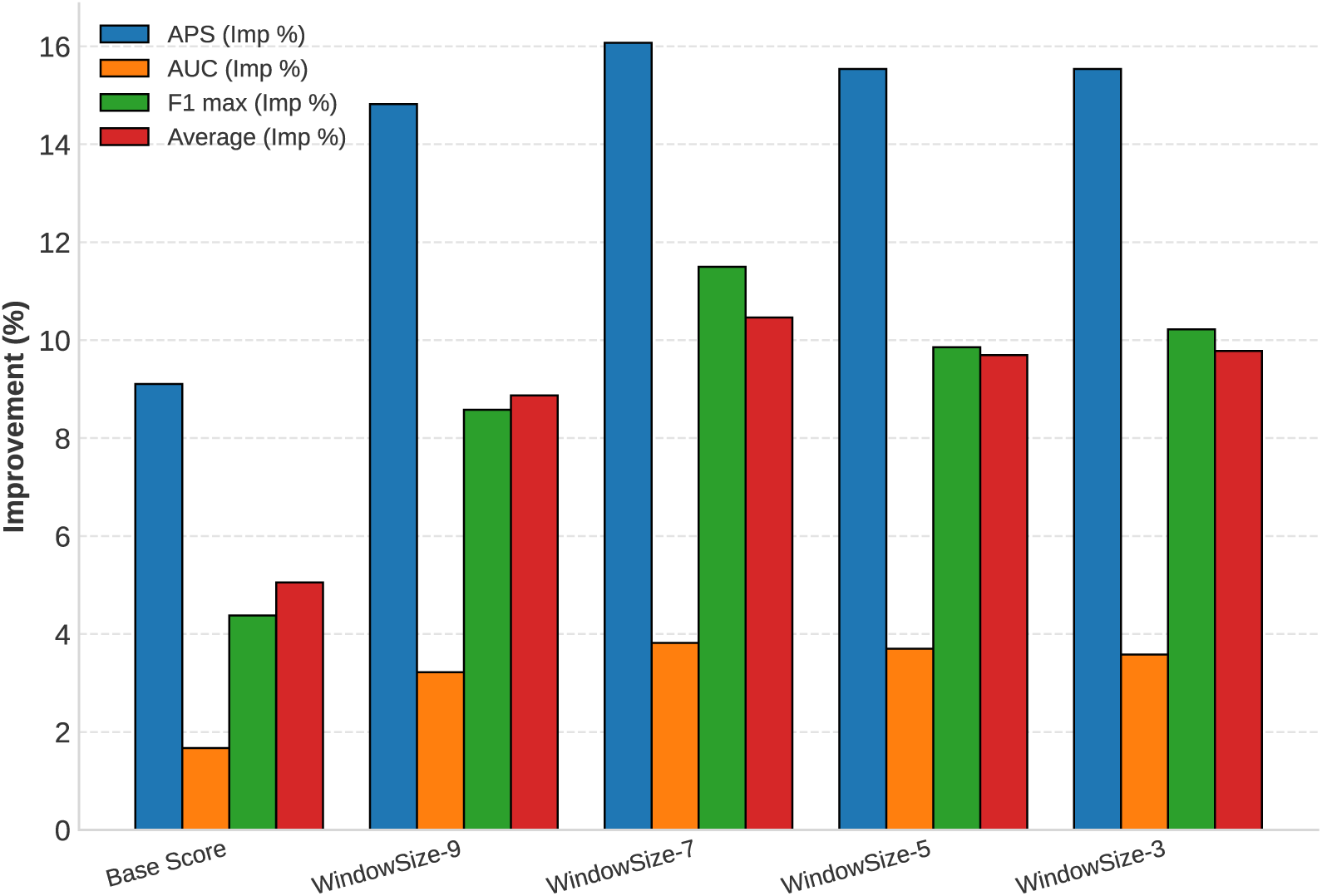
Impact of varying window sizes for ESM2-derived statistical features on model performance metrics. Percentage improvements are compared across different window sizes (3, 5, 7, 9), and window size 7 yields the best overall performance.

#### 1.3.6 Effect of Smoothing Techniques

To further reduce local fluctuations in the predicted probability scores, we applied several smoothing techniques. These fluctuations are most pronounced in order–disorder transition regions, where the predicted state switches between ordered and disordered segments. By smoothing the output scores, we generated more coherent disorder profiles. Figure 12 provides a residue-level plot of disorder probability predictions. The blue line shows the original raw probability values, which fluctuate considerably between residues, while the red line represents the smoothed probabilities. The smoothed curve offers a clearer and more interpretable trend, highlighting regions with high disorder probability (approaching 1.0) and areas of low disorder likelihood (closer to 0.2). This visual comparison confirms that smoothing helps identify coherent disorder patterns within protein sequences.

**Figure 12.**
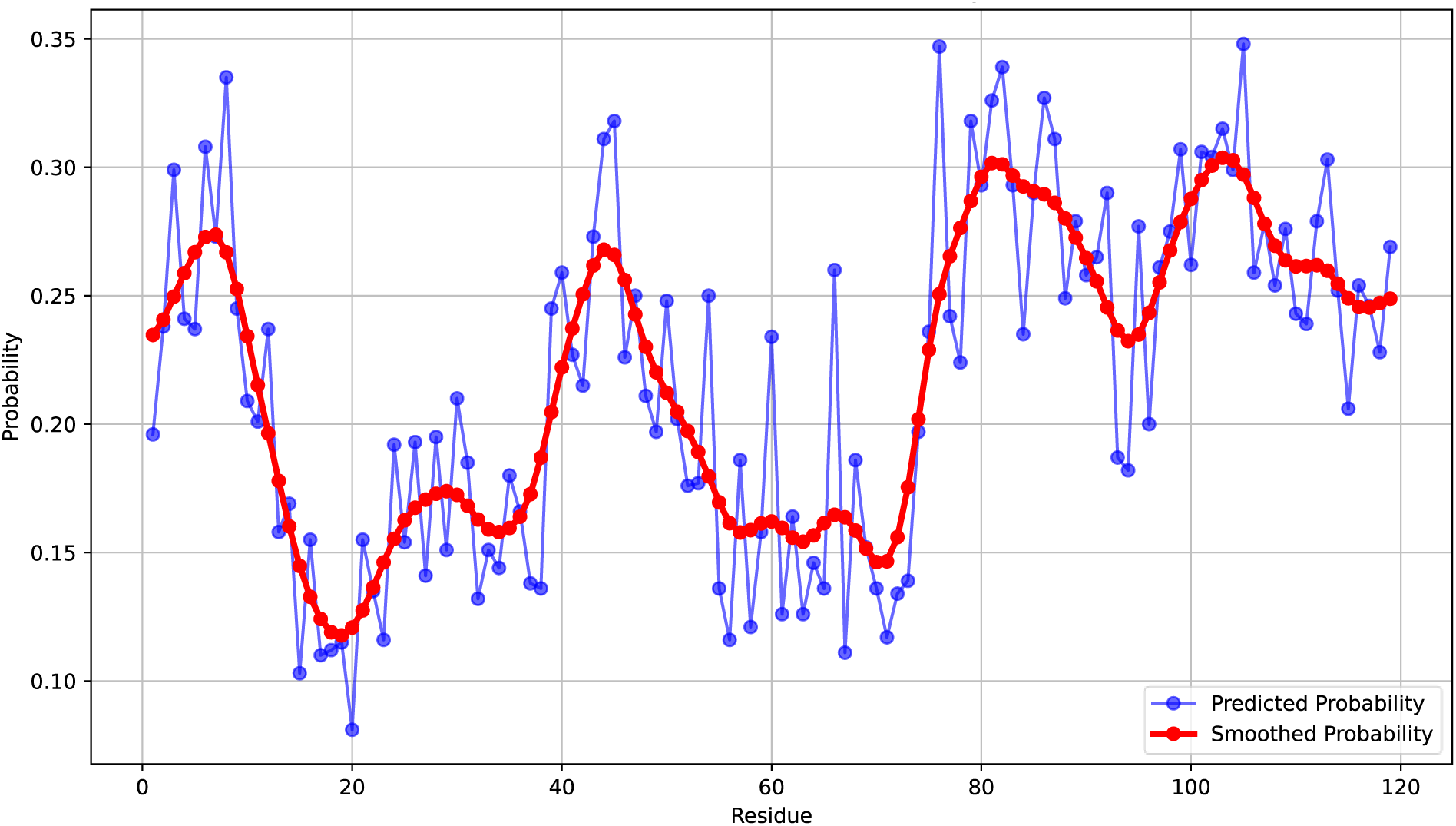
The effect of smoothing in disordered proteins. The *x*-axis represents the residue position, while the *y*-axis shows the probability of disorder. The blue line represents the original probability predictions for each residue, displaying considerable fluctuations. The red line represents the smoothed probability values, offering a clearer trend. Overall, smoothing suppresses residue-to-residue noise and produces a more reliable disorder probability.

We evaluated the effects of moving average, exponential moving average, and Gaussian smoothing methods. As illustrated in Figure 13, all three smoothing techniques led to incremental improvements in ROC AUC performance when compared to the baseline (unsmoothed) predictions. The Gaussian smoothing method achieved the best results, followed closely by the exponential moving average, indicating their effectiveness in capturing broader contextual patterns without overly distorting localized predictions. These results suggest that incorporating smoothing as a post-processing step can enhance the performance of disorder prediction models.

**Figure 13.**
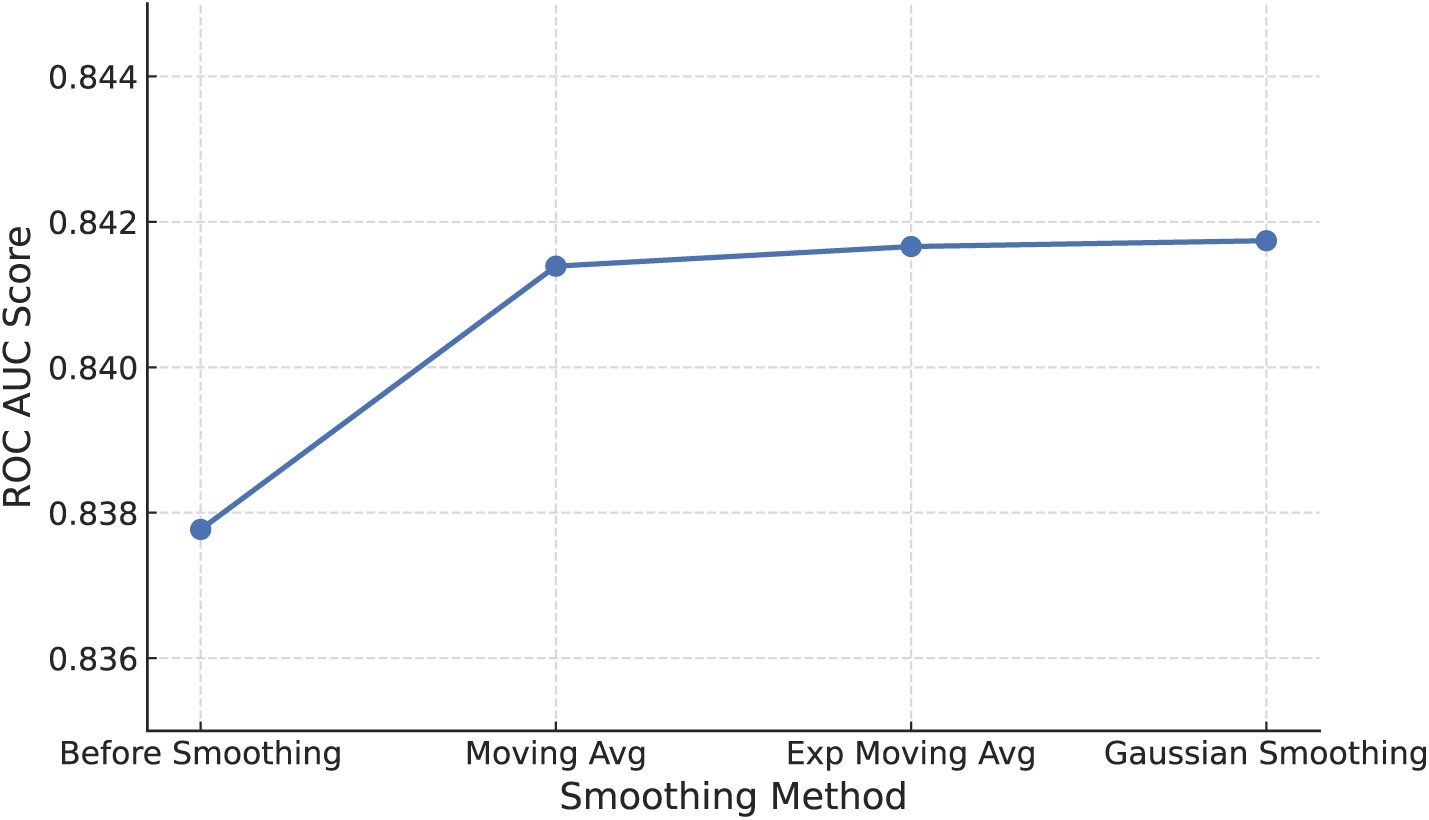
Impact of different smoothing techniques on ROC AUC performance. The ROC AUC scores before and after applying moving average, exponential moving average, and Gaussian smoothing methods are compared. The figure shows that the smoothing provides consistent improvements in ROC–AUC.

### 1.4 Architecture of ESMDisPred

We developed and trained four model variants (Figure 14) within the ESMDisPred framework, focusing on different feature combinations, sequence representations, and algorithms. Table 1 summarizes the variants and their key differences. Three ESMDisPred-LightGBM variants were submitted to the CAID3 challenge, each designed to evaluate the contribution of distinct feature sets. ESMDisPred-1 utilized features from DisPredict3.0 and ESM1, ESMDisPred-2 incorporated both DisPredict3.0 and ESM1 features along with additional ESM2-derived representations, and ESMDisPred-2PDB further extended the configuration by removing missing residues based on experimental structures. Across multiple benchmark datasets in the CAID3 challenge, these models consistently outperformed existing state-of-the-art tools in terms of ROC AUC, AP, and F1 max.

**Table 1.** Summary of ESMDisPred model variants evaluated in this study, including the learning algorithm, input feature sets/representations, and preprocessing differences.

| Variant | Algorithm / Architecture | Feature / Representation Set | Submitted to CAID3? | Purpose |
| --- | --- | --- | --- | --- |
| ESMDisPred-1 | LightGBM | DisPredict3.0 + ESM1 | Yes | Baseline feature set (DisPredict3.0 + ESM1) |
| ESMDisPred-2 | LightGBM | DisPredict3.0 + ESM1 + ESM2 | Yes | Added value of ESM2 representations |
| ESMDisPred-2PDB | LightGBM | DisPredict3.0 + ESM1 + ESM2 | Yes | Effect of structure-aware residue filtering |
| ESMDisPred-DNN | Transformer-CNN deep neural network | Sequence representations (same input features as ESMDisPred-2PDB pipeline) | No | Assesses whether a hybrid CNN (local motifs) + Transformer (long-range context) architecture improves per-residue disorder prediction |

**Figure 14.**
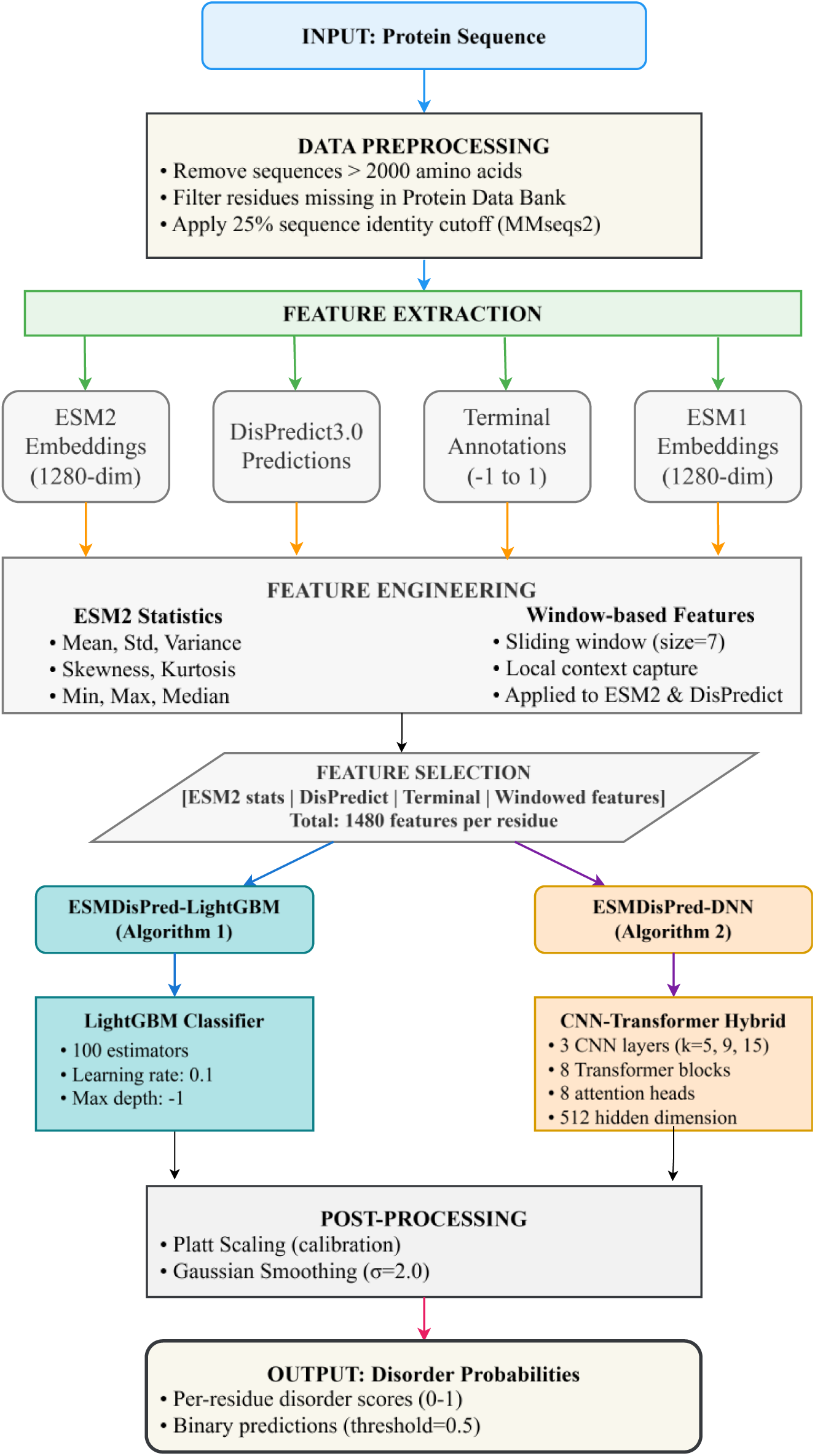
ESMDisPred pipeline for protein disorder prediction. The workflow begins with protein sequence preprocessing. Feature extraction combines four sources: ESM2 embeddings, DisPredict3.0 predictions, terminal annotations for N/C-terminal residues, and ESM1 embeddings. Feature engineering applies statistical transformations to ESM2 embeddings and generates windowed features from neighboring residues. The resulting features per residue are processed by two model variants: ESMDisPred-LightGBM and ESMDisPred-DNN (CNN-Transformer architecture). Both models produce binary disorder predictions and probability scores for each residue.

The first three variants share an identical architectural design but differ in their input feature compositions. The ESMDisPred-LightGBM (Algorithm 1), architecture integrates pretrained protein language model embeddings with a LightGBM classifier to predict intrinsically disordered regions in protein sequences. The process begins with preprocessing protein sequences to remove long or redundant entries. For each sequence, embeddings are generated using the ESM2 language model, which captures contextual and evolutionary features from raw amino acid sequences. These embeddings are transformed into statistical features such as mean, variance, and kurtosis and enriched with local context through sliding window techniques. Additionally, disorder-specific features are extracted using the DisPredict3.0 tool. The resulting features are then used to train a LightGBM model, which is optimized using cross-entropy loss and tuned hyperparameters (e.g., number of estimators). During inference, features from both ESM2 and DisPredict3.0 are computed for test sequences and passed into the trained LightGBM model to generate residue-level disorder probabilities. Finally, smoothing is applied to refine the output. This architecture combines the representational power of deep protein embeddings with the gradient-boosted trees, resulting in a high-performing disorder prediction system.

#### Algorithm 1

**ESMDisPred – Disorder Prediction using LightGBM (ESMDisPred-LightGBM)**

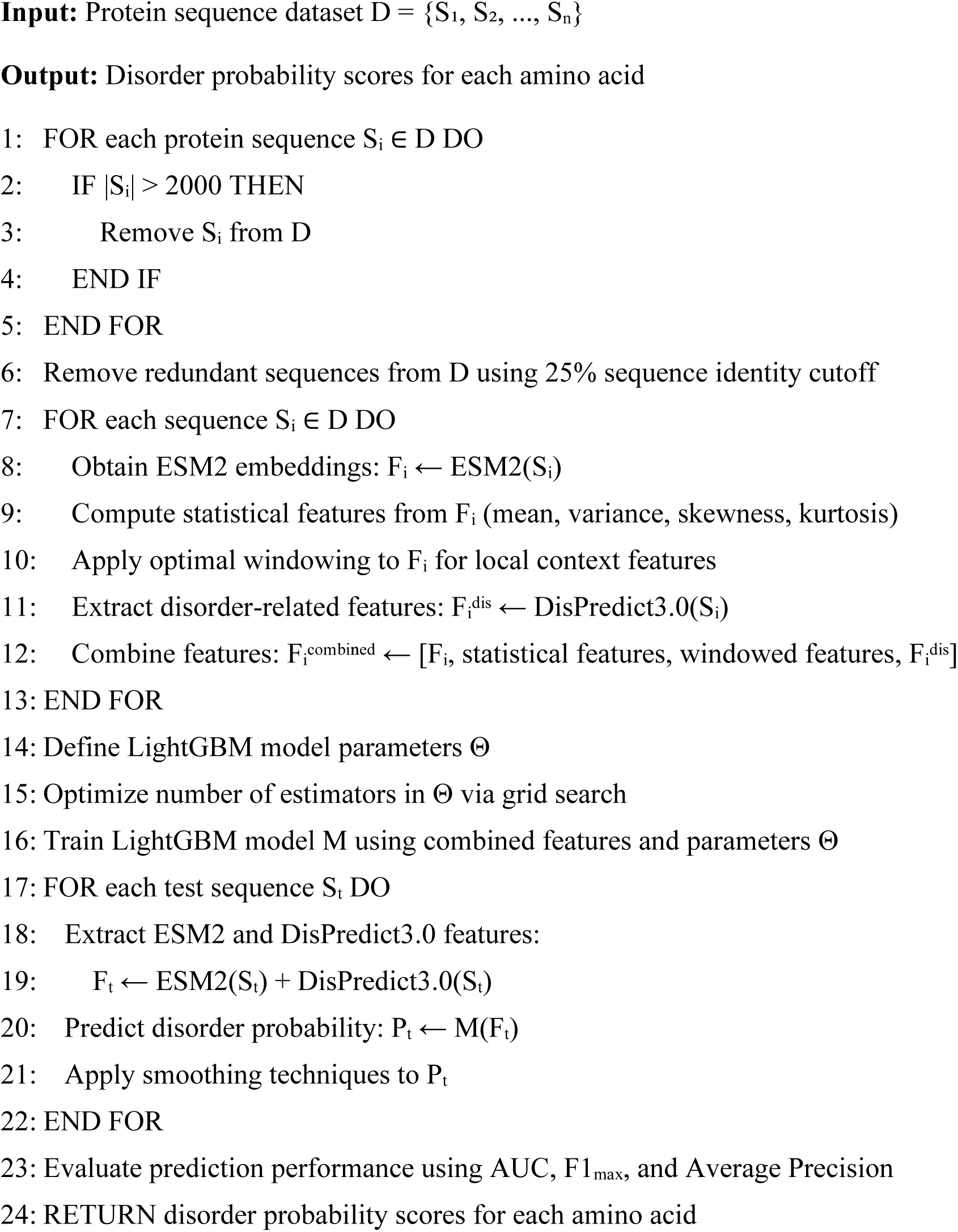

To further enhance the predictive performance of IDP after the CAID3 challenge, we designed ESMDisPred-DNN, an advanced Transformer-CNN-based deep neural network. As outlined in Algorithm 2, this model employs a hybrid architecture that integrates convolutional neural networks (CNNs) and Transformer encoders to capture both local and long-range dependencies within protein sequences. The motivation for this design arises from the observation that disordered proteins often contain both short and long disordered regions, requiring a model capable of learning across multiple spatial scales. The CNN layers effectively identify local sequence motifs, while the Transformer layers capture contextual relationships over longer sequence spans, making the architecture particularly well-suited for per-residue disorder prediction.

In ESMDisPred-DNN, we formulate residue-level IDR prediction as a per-token binary classification problem over variable-length protein sequences. For a given sequence of length *L* each residue *t* ∈ 1,2,…, *L* is represented by a standardized feature vector 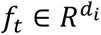. All features are z-score normalized using statistics computed from the training split and subsequently applied to the validation and test sets to ensure consistent scaling. During training, mini-batches are padded to the length of the longest sequence within each batch. A key-padding mask *M* is applied to ensure padded positions are excluded from both attention computation and loss calculation, preventing them from affecting model predictions or training.

Each residue feature is projected to the model dimension and combined with fixed sinusoidal positional encodings (see Eq. (5)):

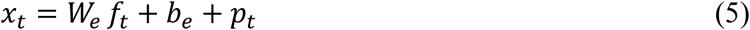

where *x*_*t*_ represents the input at position *t* in the model’s embedding space, 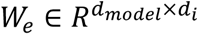 and 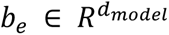 are learnable linear projections that map the input features from dimension *d*_*i*_ to the model’s embedding dimension *d_model_*. 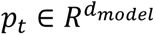 represents the fixed sinusoidal positional encoding.

Before global attention, a lightweight, length-preserving convolutional stem injects local inductive bias via three depthwise 1D convolutions (kernels 5, 9, 15), each followed by a pointwise projection. We added convolutional layer so that the model can learn the short- and mid-range motifs. Sequence context is integrated by an eight-block pre-norm Transformer (LayerNorm → sublayer → residual). Within each block, queries (*Q*), keys (*K*), and values (*V*) are formed from the hidden states 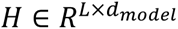 as shown in Eq. (6).

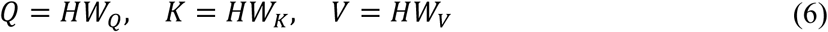

Where *W*_*Q*_, *W*_*K*_, 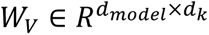 are linear projection matrices produce the queries, keys, and values. Masked multi-head self-attention is then computed as follows (see Eq. (7)):

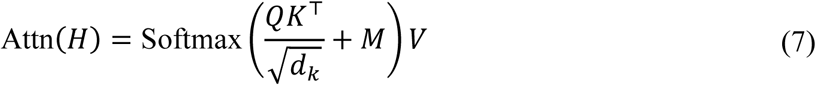

where *d*_*k*_ is the per-head key/query size and *M* injects the padding mask into the attention logits. Using pre-norm residual structure, block *l* updates are (see Eq. (8)):

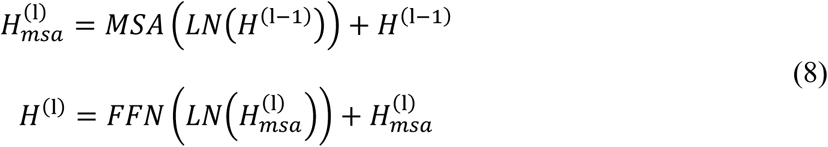

where *LN* is LayerNorm, Multi-Head Self-Attention (MSA) concatenates attention heads. The output of this layer is a position-wise feed-forward network (FFN) (see Eq. (9)). We chose GELU as the activation function because it often outperforms ReLU and tanh in Transformer architectures (e.g., BERT, GPT, ViT) [48, 49]. GELU also better models nonlinearity in continuous feature spaces, especially for embeddings or attention outputs that are approximately Gaussian-distributed.

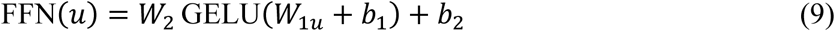

For residue-wise classification, we apply dropout to the final hidden state *h*_*t*_ and a position-wise linear layer to obtain a logit *z*_*t*_ and a disorder probability 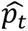 (see Eq. (10)):

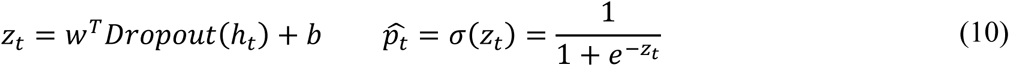

During training, we minimize a masked binary cross-entropy loss computed only over valid (non-padded) residues. We use macro per-protein ROC–AUC which is computed by averaging AUC values across proteins in each epoch. Early stopping monitors this macro-AUC, and the best-performing checkpoint is restored after training. To improve calibration, the output probabilities are refined via Platt scaling [50] (see Eq. (11)), where the calibrated probabilities are:

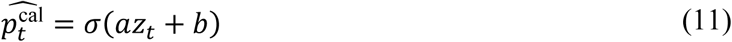

where *a and b* are learned by logistic regression. The calibrated probabilities are subsequently smoothed along the sequence using a Gaussian kernel with bandwidth σ_*g*_ > 0 and normalization constant *Z* > 0 (see Eq. (12)):

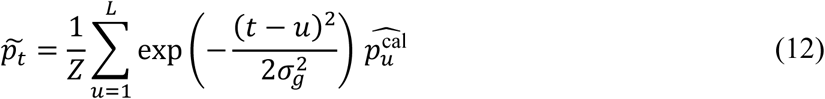

Here, *Z* ensures proper normalization of the Gaussian weights, *t* denotes the target residue position, and *u* iterates over all sequence positions contributing to the smoothed estimate. The smoothing bandwidth σ_*g*_ controls how broadly neighboring residues influence the prediction.

This architectural choice balances model complexity, training stability, and computational efficiency, which are essential for accurate residue-level predictions in long protein sequences. By combining a multi-kernel convolutional stem with Transformer-based global attention, the model effectively captures both local sequence motifs and long-range dependencies. Details of the model’s optimal hyperparameter settings are discussed in the Hyperparameter Selection section. Figure 15 shows the ESMDisPred-DNN architecture.

**Figure 15.**
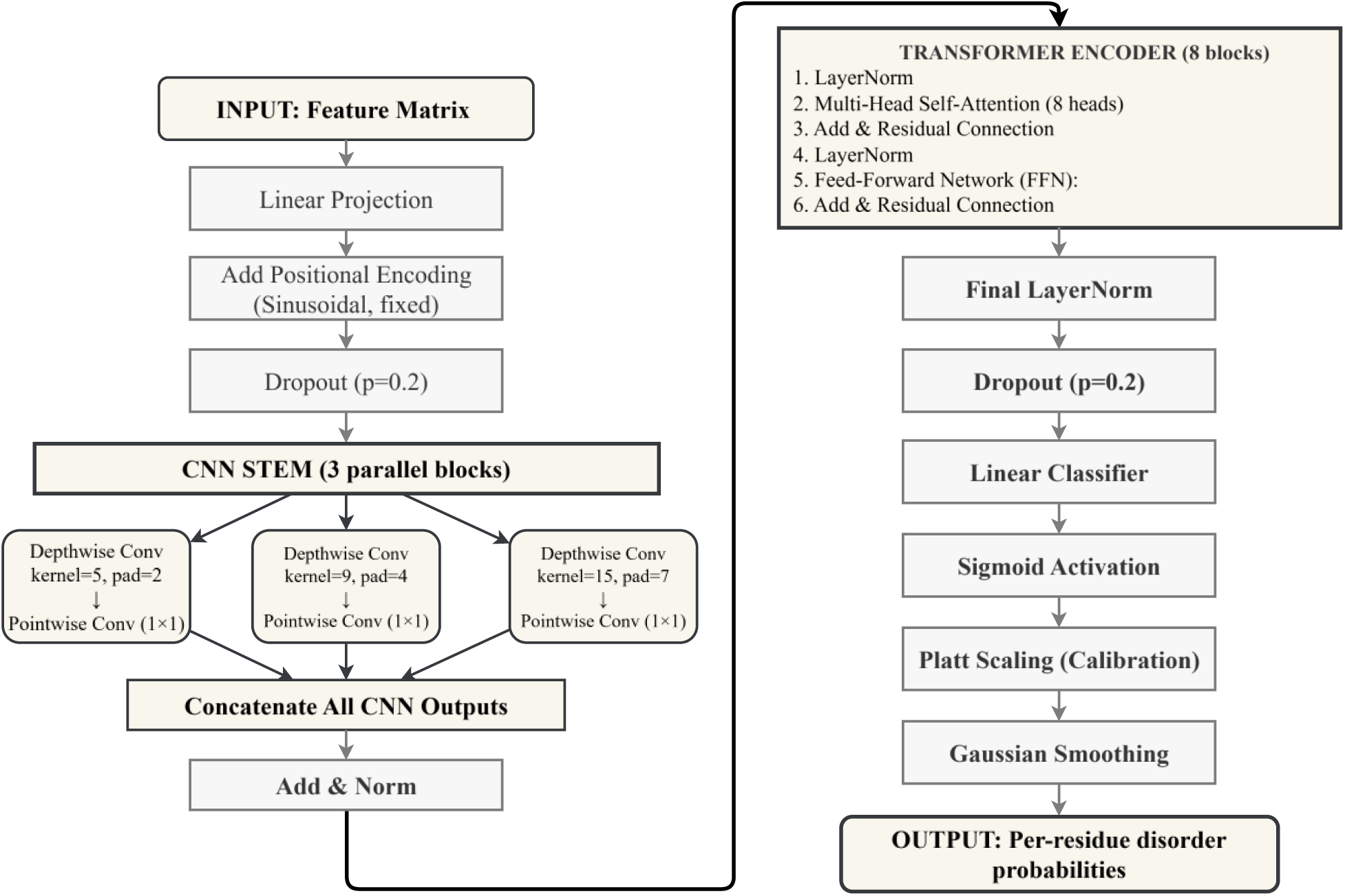
Neural network architecture of ESMDisPred-DNN for disorder prediction. The model consists of three main components: (1) Input processing with linear projection and sinusoidal positional encoding, (2) CNN stem with three parallel depthwise separable convolution blocks (kernel sizes 5, 9, and 15) that capture multi-scale local patterns, concatenated and processed through Add & Norm layers, and (3) Transformer encoder with 8 blocks, each containing multi-head self-attention (8 heads), layer normalization, feed-forward networks, and residual connections. The output passes through final layer normalization, dropout (p = 0.2), and a linear classifier to produce per-residue disorder predictions.

#### Algorithm 2

**Transformer-based Per-Residue Disorder Prediction (ESMDisPred-DNN)**

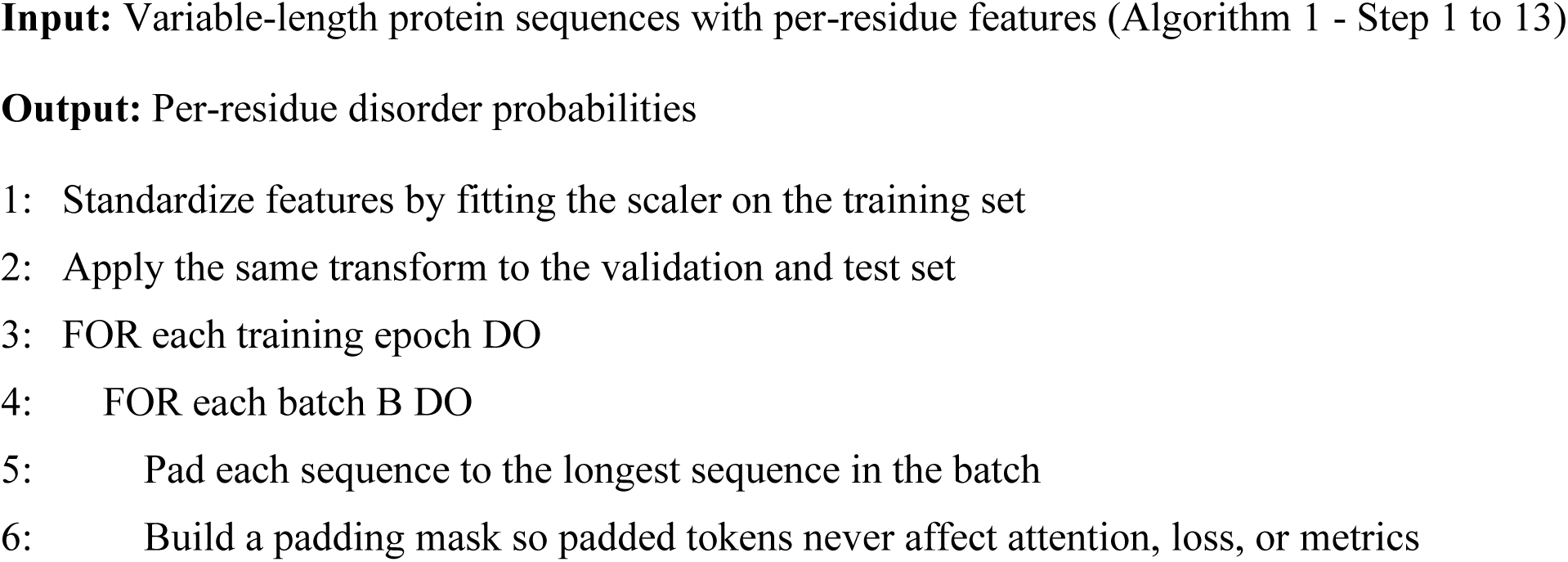
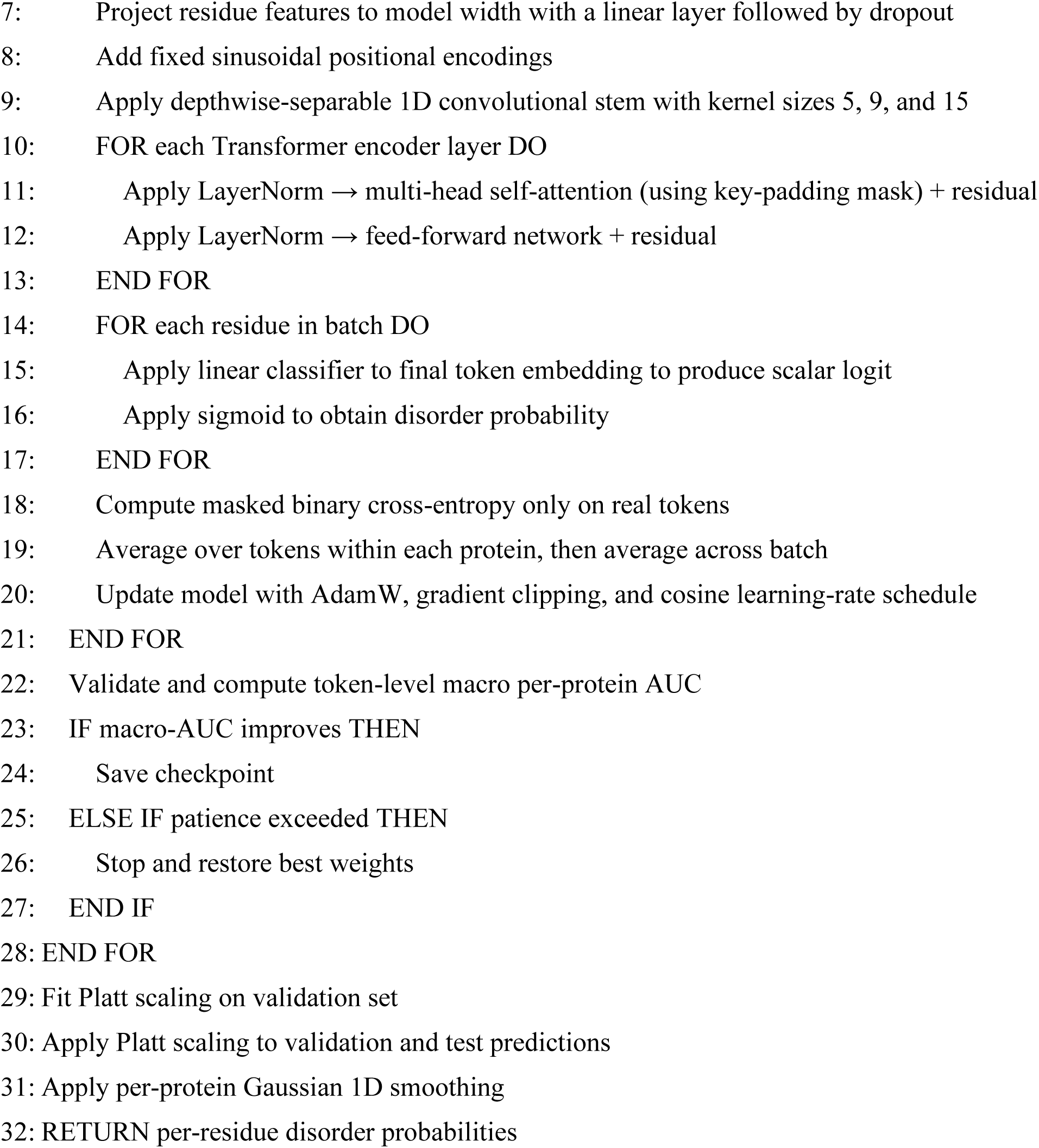

### 1.5 Experimental Results

This section presents experimental results evaluating ESMDisPred on CAID benchmarks. We first compare multiple machine learning algorithms on the validation set to select the optimal approach. Next, we analyze the enhanced ESMDisPred-DNN architecture and demonstrate its improvements over the LightGBM-based variant. We then benchmark all ESMDisPred variants against state-of-the-art disorder predictors using ROC and Precision-Recall curves. Statistical significance of ROC–AUC differences was assessed using DeLong’s test, a standard nonparametric method for comparing correlated AUCs computed on the same benchmark set, and consistent with CAID evaluation practice. Finally, we assess computational efficiency and runtime performance.

#### 1.5.1 Model Comparison and Selection on the Validation Set

A comparative performance evaluation of several machine learning models, such as LightGBM, CatBoost, XGBoost, Random Forest, and Logistic Regression, was conducted on the Disorder NOX dataset from the CAID2 benchmark. Figure 16 reports performance relative to the state-of-the-art (SOTA) model. Among all the methods, LightGBM shows the highest performance, achieving improvements of up to 7.5% in AP, 6.2% in AUC, 8.9% in F1 max, and 7.5% in the overall average performance.

**Figure 16.**
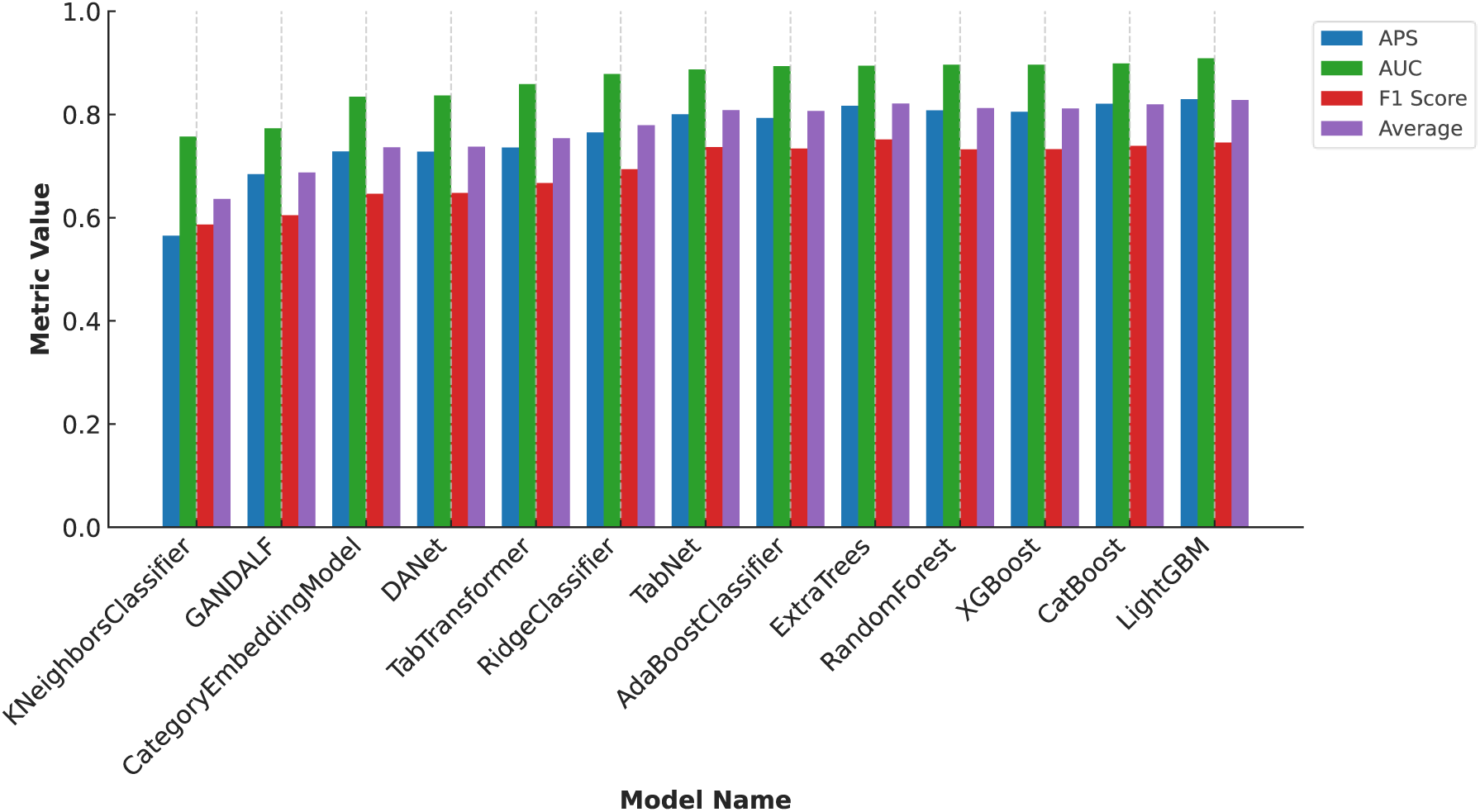
Performance comparison of various machine learning models (LightGBM, CatBoost, XGBoost, Random Forest, Logistic Regression) on the Disorder NOX Dataset (CAID2). LightGBM achieves the best overall performance across metrics and is therefore selected for the ESMDisPred-LightGBM variants.

These results demonstrate LightGBM’s superior capacity to capture complex relationships in protein sequence-derived features. Its gradient boosting framework, efficient handling of large-scale data, and robustness against overfitting make it particularly well-suited for the task of disordered region prediction. In contrast, while CatBoost and XGBoost offer competitive results, they fall short of LightGBM’s consistency across all metrics. Logistic Regression and Random Forest, being more traditional models, lag behind, highlighting the advantages of modern gradient boosting techniques.

Based on these results, we focused on optimizing the number of estimators in ESMDisPred-LightGBM, which controls the number of sequential learners in the ensemble. This parameter was chosen because of its direct influence on model complexity and predictive performance in boosting-based algorithms. Increasing the number of estimators typically reduces bias and improves accuracy, but can also increase the risk of overfitting and computational cost. Figure 17 presents the average performance improvement as a function of the number of estimators. The model achieved its highest improvement at around 100 estimators. Beyond this point, performance showed minor fluctuations and no consistent improvement, eventually declining slightly after 1500 estimators.

**Figure 17.**
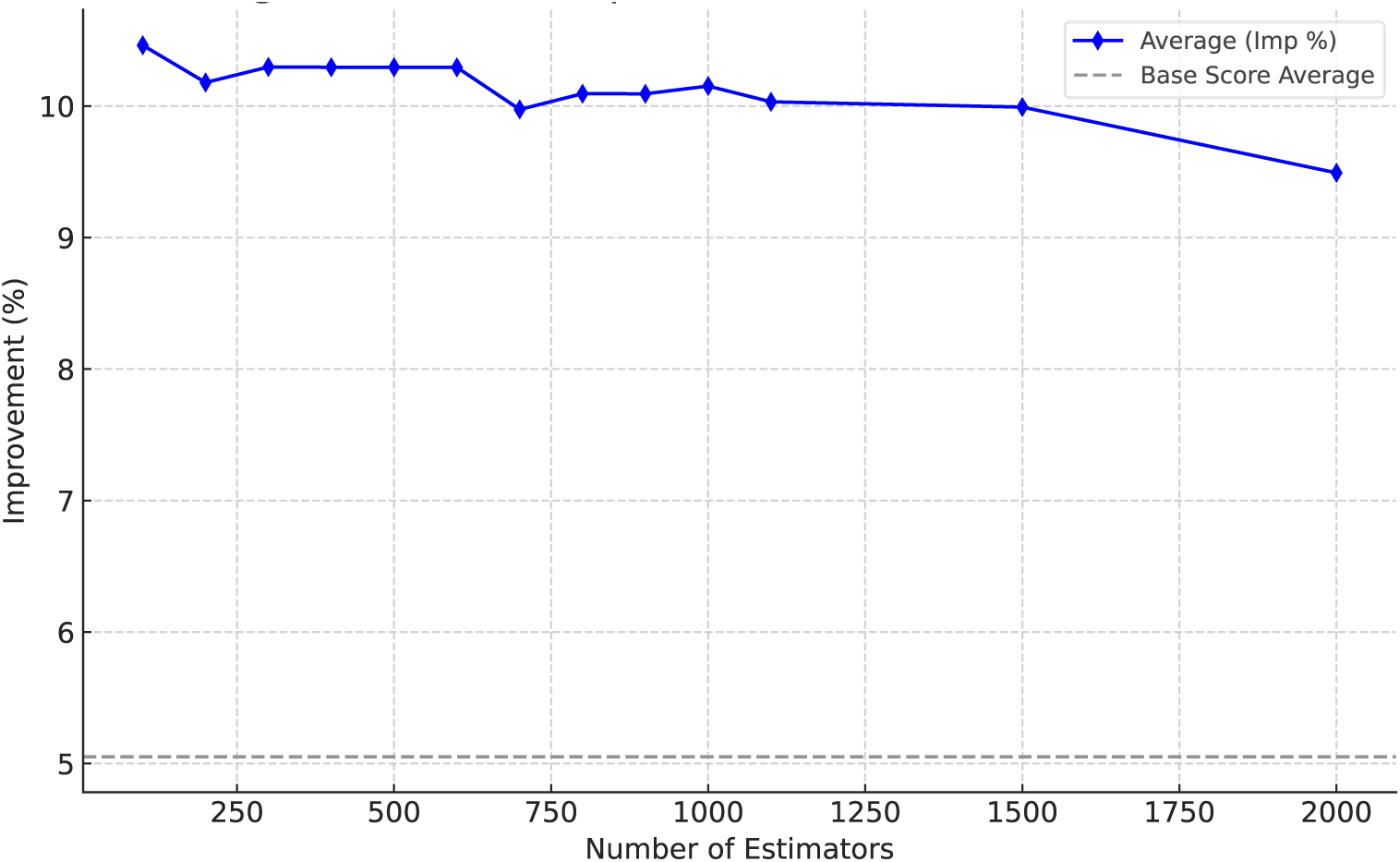
Effect of the number of estimators on LightGBM performance. The average improvement (%) is shown as the number of estimators increases, with the dashed line indicating the baseline average score. Performance is relatively stable across a wide range of estimator counts, with only marginal gains beyond a few hundred trees.

For the ESMDisPred-DNN model, we explored hyperparameters to understand how capacity, regularization, and post-hoc calibration affect generalization and performance. A grid search was conducted over the ranges of parameters (Table 2): projection width, attention heads, Transformer depth, feed-forward width, dropout rates, weight decay, and learning rates. Each model was trained with early stopping based on the macro AUC metric. We first calculated the AUC for each protein sequence individually, based on residue-level disorder probabilities, and then averaged these values across all proteins. This per-protein macro averaging ensures that each protein contributes equally to the evaluation, regardless of sequence length or class imbalance. Such an approach prevents bias towards majority-class (ordered) sequences.

**Table 2.** Hyperparameter search space with minimum and maximum values, along with the number of unique sampled values used during model optimization.

| **Hyperparameter** | **Min** | **Max** | **Unique values** |
| --- | --- | --- | --- |
| Projection Dimension | 256 | 512 | 4 |
| Number of Attention Heads | 8 | 16 | 4 |
| Number of Layers | 4 | 8 | 3 |
| Feed-Forward Dimension | 768 | 2560 | 10 |
| Dropout Rate | 0.2 | 0.3 | 3 |
| Learning Rate | 0.0001 | 0.0003 | 3 |
| Weight Decay | 0.01 | 0.05 | 3 |

Across 35 independent training runs (Table 3), model performance consistently clustered within a narrow range, indicating that the prediction task is robust to moderate architectural and regularization variations. Table 2 summarizes all 35 ESMDisPred-DNN configurations sampled during the grid search, along with their corresponding validation AUC values.

**Table 3.**
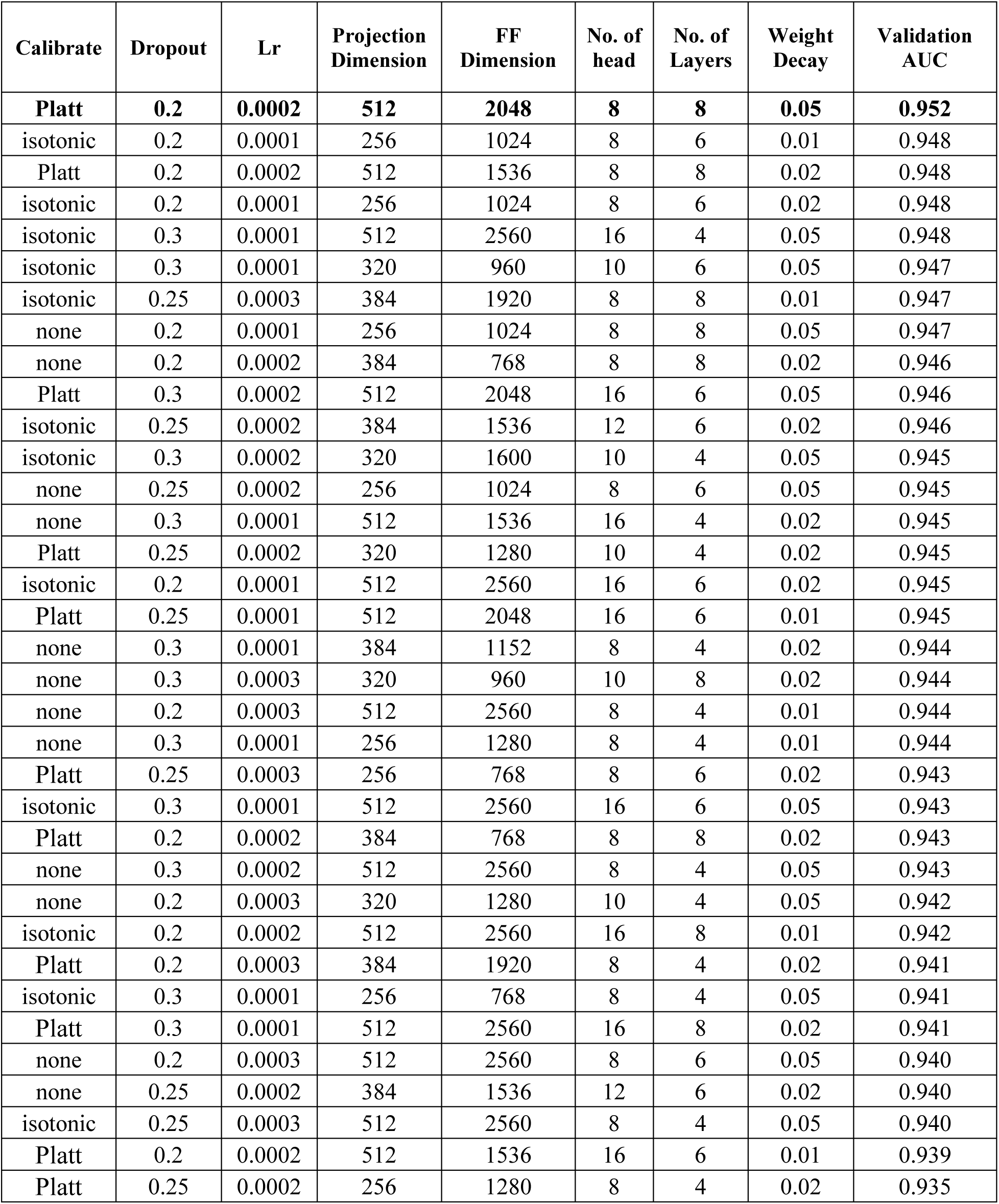
Summary of all 35 ESMDisPred-DNN training runs. Each row represents a unique hyperparameter configuration sampled during the grid search, with corresponding validation performance metrics. Boldface indicates the best-performing hyperparameter configuration (highest validation AUC), which was selected for the ESMDisPred-DNN model.

Based on the experimental results, the selected model configuration for validation was a 512-dimensional, 8-head, 8-layer Transformer with feed-forward width 2048, dropout 0.2, learning rate 2×10^-4^, weight decay 0.05, and Platt calibration for our final training model. This configuration achieved a validation AUC of 0.952, along with a test AUC of 0.895, and a test Average Precision (AP) ≈ of 0.778. Figure 18 summarizes how calibration affects generalization across model configurations by plotting validation AUC against test AUC for three settings (no calibration, isotonic regression, and Platt scaling) [50, 51]. Each point corresponds to a distinct hyperparameter configuration. Overall, validation AUC tracks test AUC, and calibration produces a modest but consistent improvement, with Platt scaling giving the strongest results across configurations.

**Figure 18.**
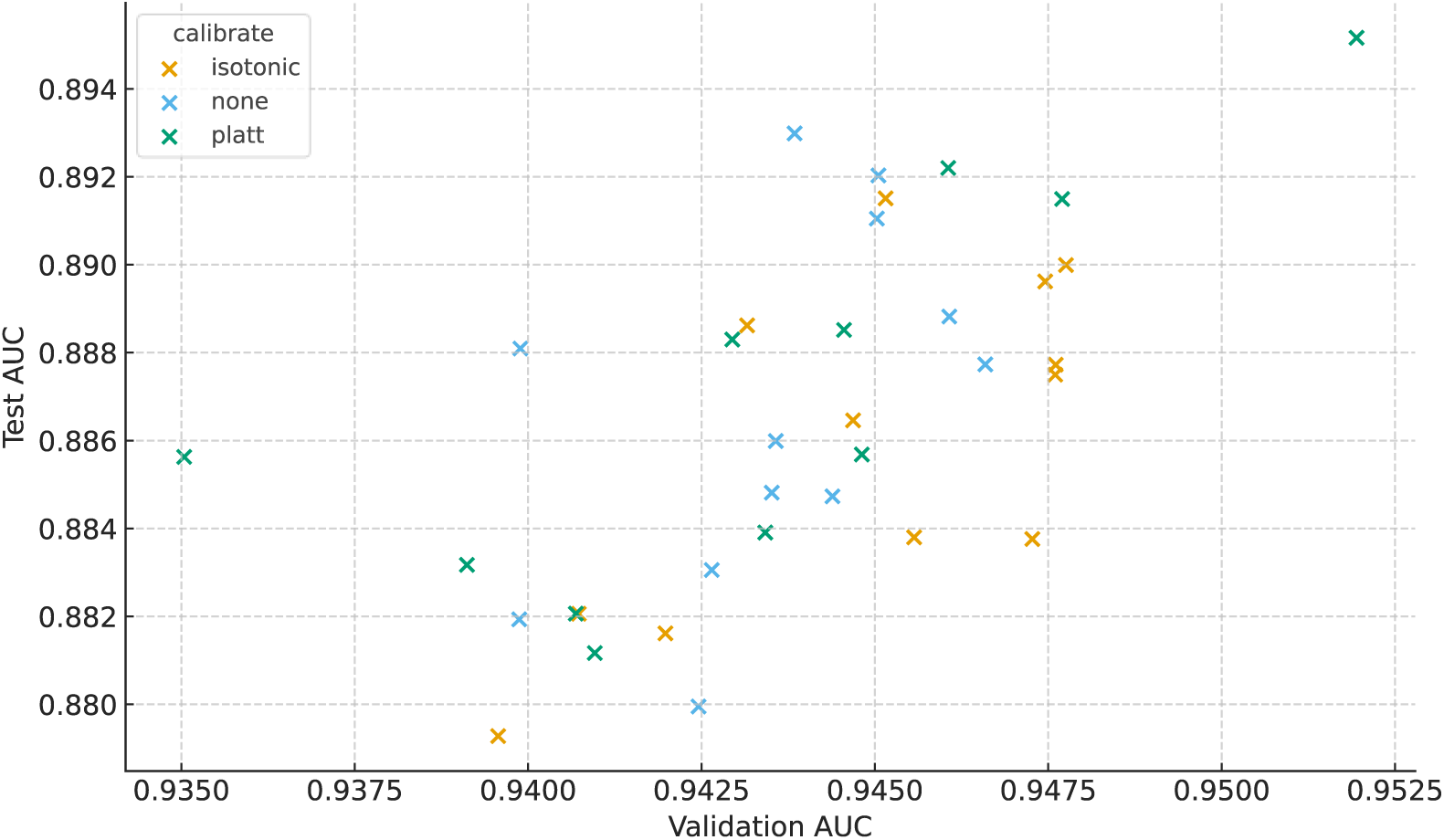
Comparison of calibration methods (none, isotonic, Platt scaling) showing the relationship between validation AUC and test AUC. Each point represents a model configuration, highlighting how calibration impacts generalization performance. The validation AUC correlates with test AUC and Platt scaling, yielding the strongest overall results.

Figure 19 illustrates the comparative performance of the ESMDisPred-DNN and ESMDisPred-LightGBM models. The results clearly demonstrate the superior performance of the ESMDisPred-DNN architecture. Unlike the LightGBM variant, ESMDisPred-DNN employs a CNN– Transformer hybrid model, as described in Section 1.4. This design leads to significant improvements in predictive accuracy.

**Figure 19.**
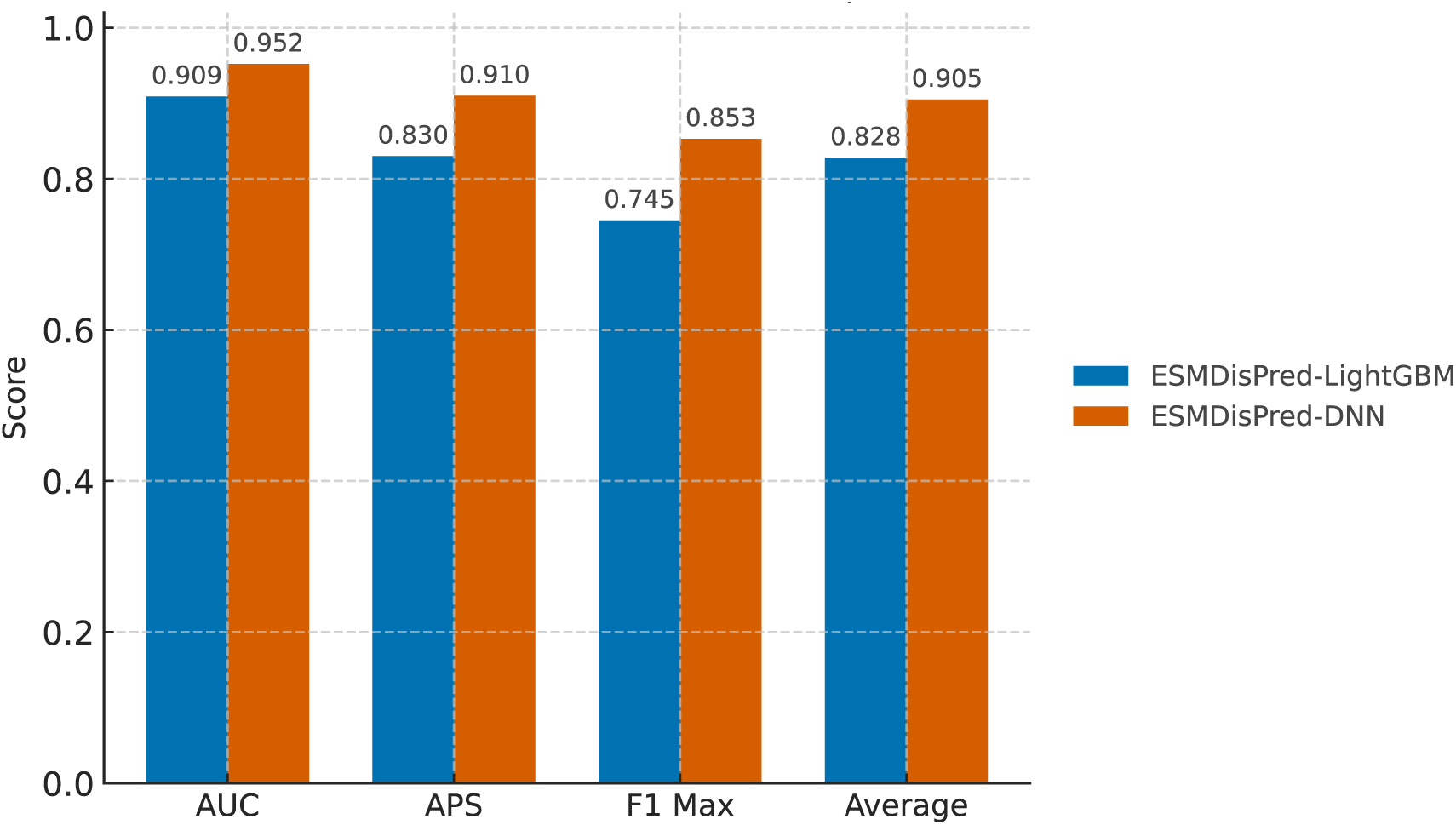
Comparison of predictive performance between ESMDisPred-LightGBM and ESMDisPred-DNN across three evaluation metrics. ESMDisPred-DNN consistently outperforms the LightGBM variant across all evaluation metrics.

#### 1.5.2 Comparison with Existing Methods

To assess the performance of ESMDisPred, we compared it with several state-of-the-art disorder prediction tools. Figure 20 and Table 3 provide a comparative evaluation of ESMDisPred models against other top-performing predictors in CAID3 challenges. The top panel of Figure 20 presents the Receiver Operating Characteristic (ROC) curves, where ESMDisPred-DNN achieves the highest AUC (0.8952), surpassing all other methods, including ESMDisPred-2PDB (AUC = 0.8855), ESMDisPred-1 (AUC = 0.8763), and ESMDisPred-2 (AUC = 0.8723). The bottom panel shows the Precision–Recall (PR) curves, where ESMDisPred-DNN again leads with a PR AUC of 0.7778, outperforming ESMDisPred-2PDB (0.7542), ESMDisPred-1 (0.7450), ESMDisPred-2 (0.7431), and other state-of-the-art predictors. These results confirm not only the discrimination ability of the ESMDisPred-LightGBM family but also the clear superiority of ESMDisPred-DNN. Importantly, ESMDisPred-DNN is a new model developed after CAID. While it was not submitted to the challenge, its performance on the validation and independent test set demonstrates that it extends the strengths of the feature-based LightGBM models with a more expressive CNN-Transformer hybrid architecture.

**Table 4.** Performance comparison of protein intrinsically disordered region (IDR) predictors on the test set, reported as AUC, average precision, and maximum F1 (higher is better). ESMDisPred-DNN attains the best overall performance.

| Methods | AUC | Average Precision | F1 max |
| --- | --- | --- | --- |
| ESMDisPred-DNN | **0.895** | **0.778** | **0.759** |
| ESMDisPred-2PDB | 0.885 | 0.754 | 0.749 |
| ESMDisPred-1 | 0.876 | 0.745 | 0.720 |
| ESMDisPred-2 | 0.872 | 0.743 | 0.724 |
| DisoFLAG-IDR | 0.868 | 0.714 | 0.671 |
| DisorderUnetLM | 0.862 | 0.702 | 0.647 |
| fIDPnn3a | 0.860 | 0.700 | 0.668 |
| UdonPred-combined | 0.854 | 0.668 | 0.671 |
| DisPredict3.0 | 0.850 | 0.666 | 0.658 |
| rawMSA | 0.850 | 0.671 | 0.641 |

**Boldface** indicates the best score in each column.

**Figure 20.**
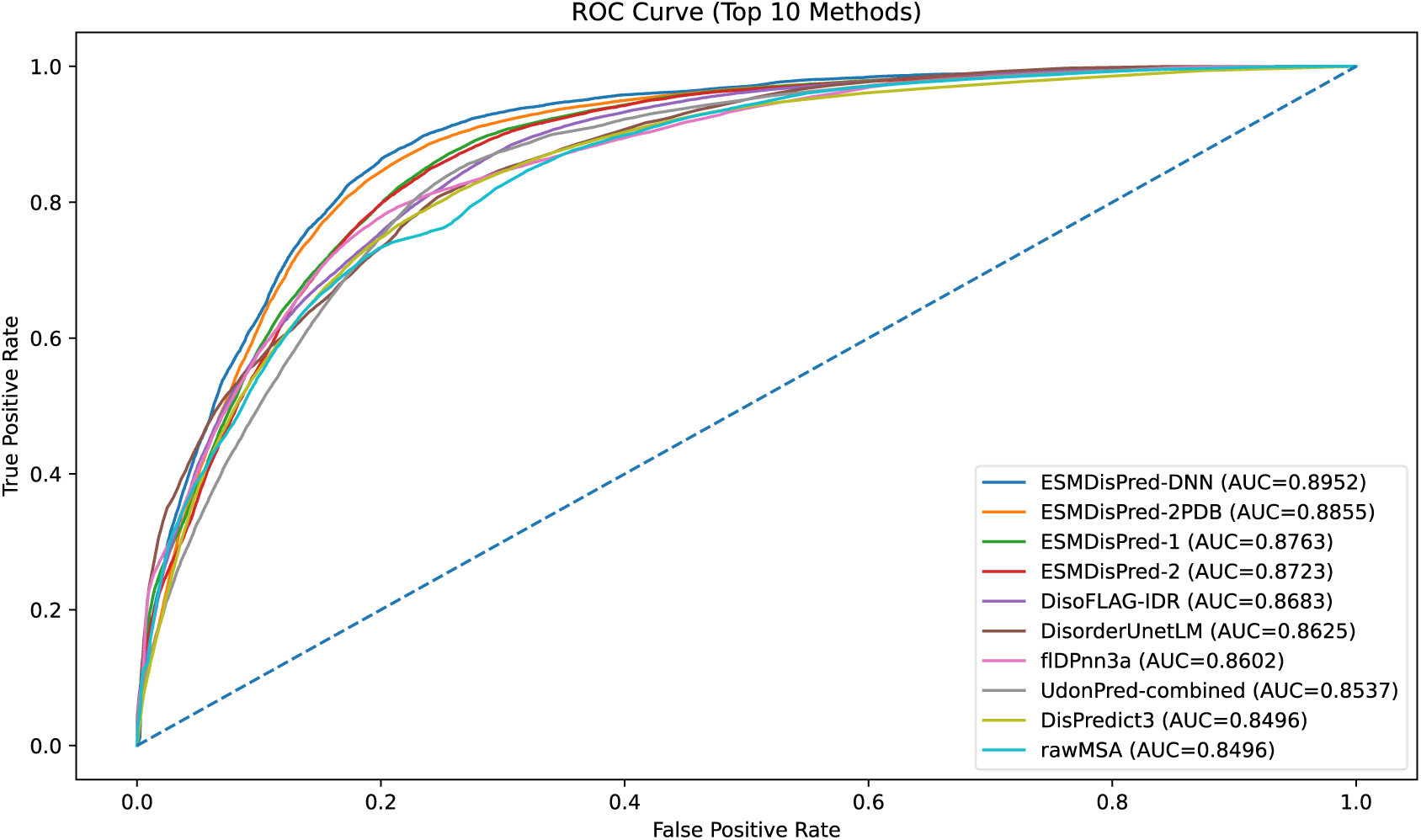
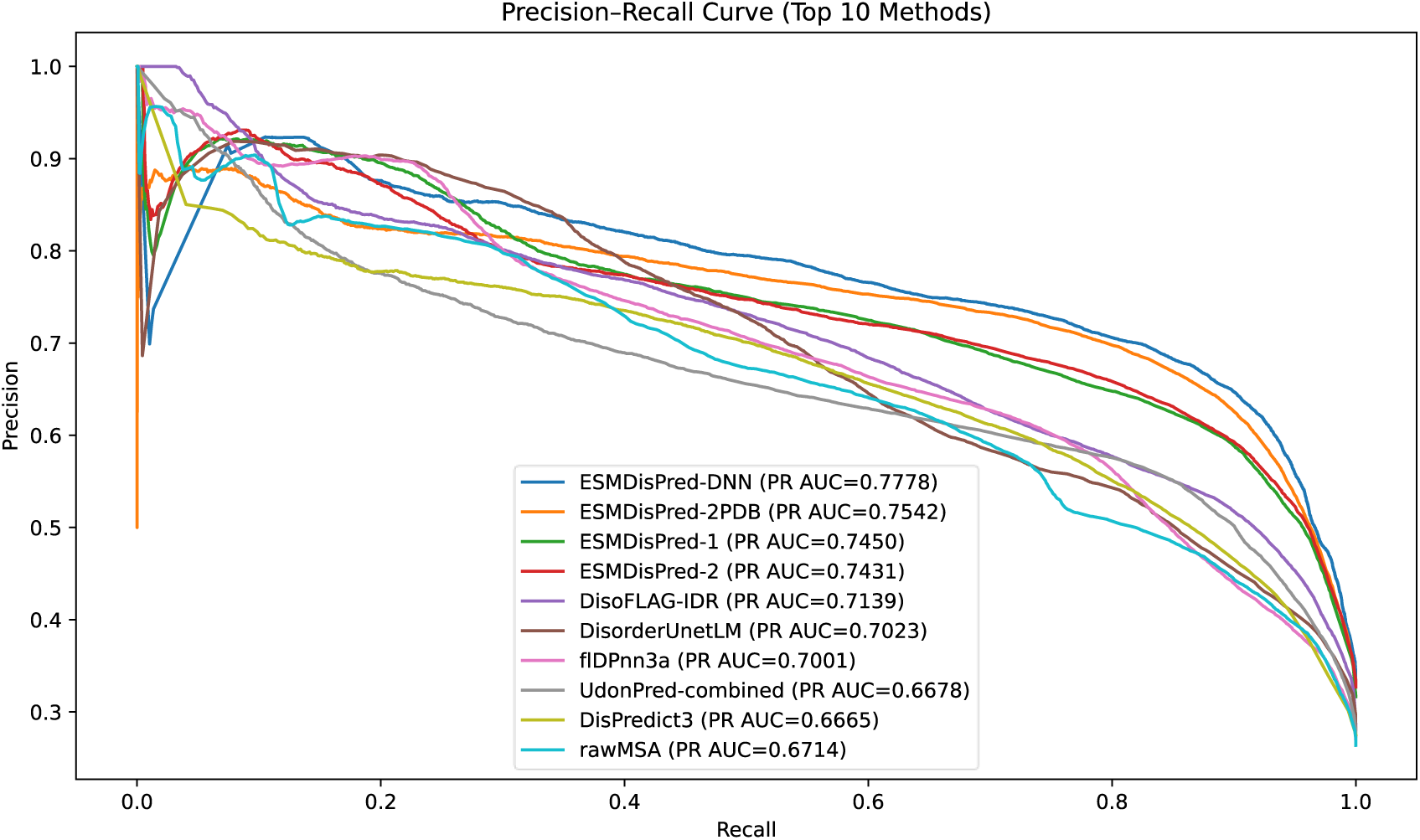
Comparison of ESMDisPred models against other top-performing predictors on the Disorder NOX dataset (CAID2). (Top) Receiver Operating Characteristic (ROC) curves showing model discrimination capabilities, with Area Under Curve (AUC) values for each method. (Bottom) Precision-Recall (PR) curves comparing predictive performance in terms of precision and recall. ESMDisPred-DNN is the best overall method among the top 10, achieving the highest ROC–AUC (∼0.895) and the highest PR–AUC (∼0.778). ESMDisPred-DNN provides the strongest discrimination and the best precision–recall trade-off, outperforming both ESMDisPred variants and state-of-the-art predictors.

To further analyze whether our model’s improvement is statistically significant compared to other approaches, we performed pairwise comparisons using the DeLong test. The DeLong test is a statistical method used to compare the area under the ROC curve (AUC) between two correlated classifiers [52]. It is particularly useful in assessing whether one prediction method performs significantly better than another, based on their true positive and false positive rates. The test assumes that AUCs follow a normal distribution and computes confidence intervals and p-values to determine statistical significance.

We compared the top 10 methods using pairwise DeLong tests for correlated ROC curves and summarized the outcomes in a significance grid (Figure 21). After Benjamini–Hochberg FDR correction (α = 0.05), blue cells indicate the row method’s AUC is significantly higher than the column method, red indicates significantly lower, and white denotes no significant difference. ESMDisPred-DNN was the top performer, achieving statistically significant gains over every comparator. A second tier included ESMDisPred-2PDB and DisorderUnetLM, which were not significantly different from each other but outperformed most remaining approaches. Within the ESMDisPred family, ESMDisPred-2 exceeded ESMDisPred-1, while DisoFLAG-IDR did not differ significantly from either. flDPnn3a lagged DisorderUnetLM, and UdonPred-combined and DisPredict3 were largely indistinguishable yet underperformed relative to the leaders. The rawMSA baseline was significantly worse than all other methods. Notably, all results except the ESMDisPred variants were collected from the CAID3 challenge submissions.

**Figure 21.**
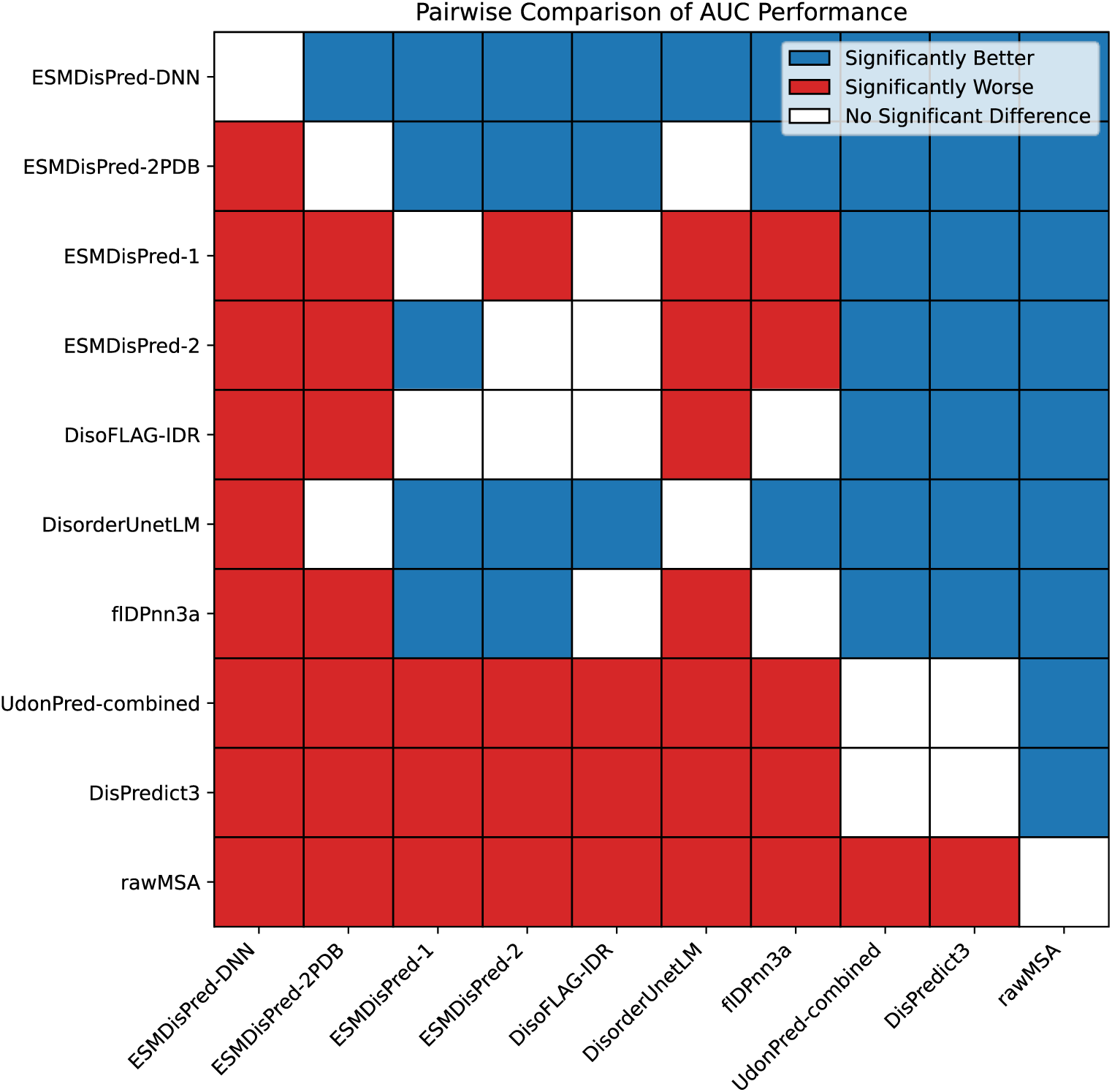
DeLong confidence interval comparison for the top 10 disorder prediction methods from the CAID (Critical Assessment of Intrinsic Disorder) results. Pairwise comparisons are based on ROC AUC scores. Blue cells indicate that the method on the row is statistically significantly better than the method on the column (p < 0.05), red indicates significantly worse performance, and white indicates no significant difference. The DeLong test was applied at the 95% confidence level using results from the official CAID challenge. ESMDisPred-DNN is statistically superior to most competing methods in ROC–AUC.

We also evaluated model performance on the Disorder-PDB dataset; however, the results did not show the same level of improvement observed on NOX. Several factors contribute to this discrepancy. First, our entire pipeline was optimized specifically for the NOX dataset, including feature engineering, hyperparameter tuning, and model architecture decisions. Second, the Disorder-PDB dataset contains proteins with experimentally resolved structures, which may exhibit different disorder characteristics compared to the NOX dataset. Proteins in PDB often have missing residues that are assumed to be disordered, but these regions may represent experimental artifacts rather than true intrinsic disorder. Third, we observed that top-performing methods on the PDB dataset predominantly incorporate AlphaFold-predicted structures or structure-derived features (such as pLDDT scores), which our current model does not utilize. These factors explain why our method, optimized for sequence-based disorder prediction on NOX, does not transfer as effectively to the structure-enriched PDB benchmark.

#### 1.5.3 Computational Complexity

ESMDisPred is designed to deliver high predictive accuracy while maintaining computational efficiency, making it suitable for high-throughput applications. All our experiments were run in containers (Docker locally and Apptainer on LONI HPC) [53] for reproducibility. We also set deterministic seeds (NumPy, PyTorch). Training and inference were performed on LONI HPC nodes with 32-core Intel Xeon Platinum 8358, 512 GB RAM, NVIDIA A100 with 40 GB memory.

When benchmarked on the CAID3 dataset comprising 232 protein sequences, ESMDisPred achieved an average runtime of ∼95 seconds per protein under the CAID3 evaluation protocol (runtime and AUC values taken from the official CAID3 results) [25]. All ESMDisPred variants have essentially the same runtime under this protocol, since they share the same ESM2 embedding step and differ only in lightweight downstream components. This efficiency is further enhanced when the model is deployed on GPU hardware, substantially reducing inference time and enabling large-scale proteomic analyses. As shown in Figure 22, ESMDisPred achieves a good balance between runtime and AUC performance, positioning itself among the top-performing predictors. Compared to other state-of-the-art models such as rawMSA or flDPnn3b, which require significantly longer runtimes, ESMDisPred maintains competitive accuracy with substantially faster execution. This favorable trade-off between speed and predictive quality underscores the utility of ESMDisPred in proteome-wide disorder prediction pipelines.

**Figure 22.**
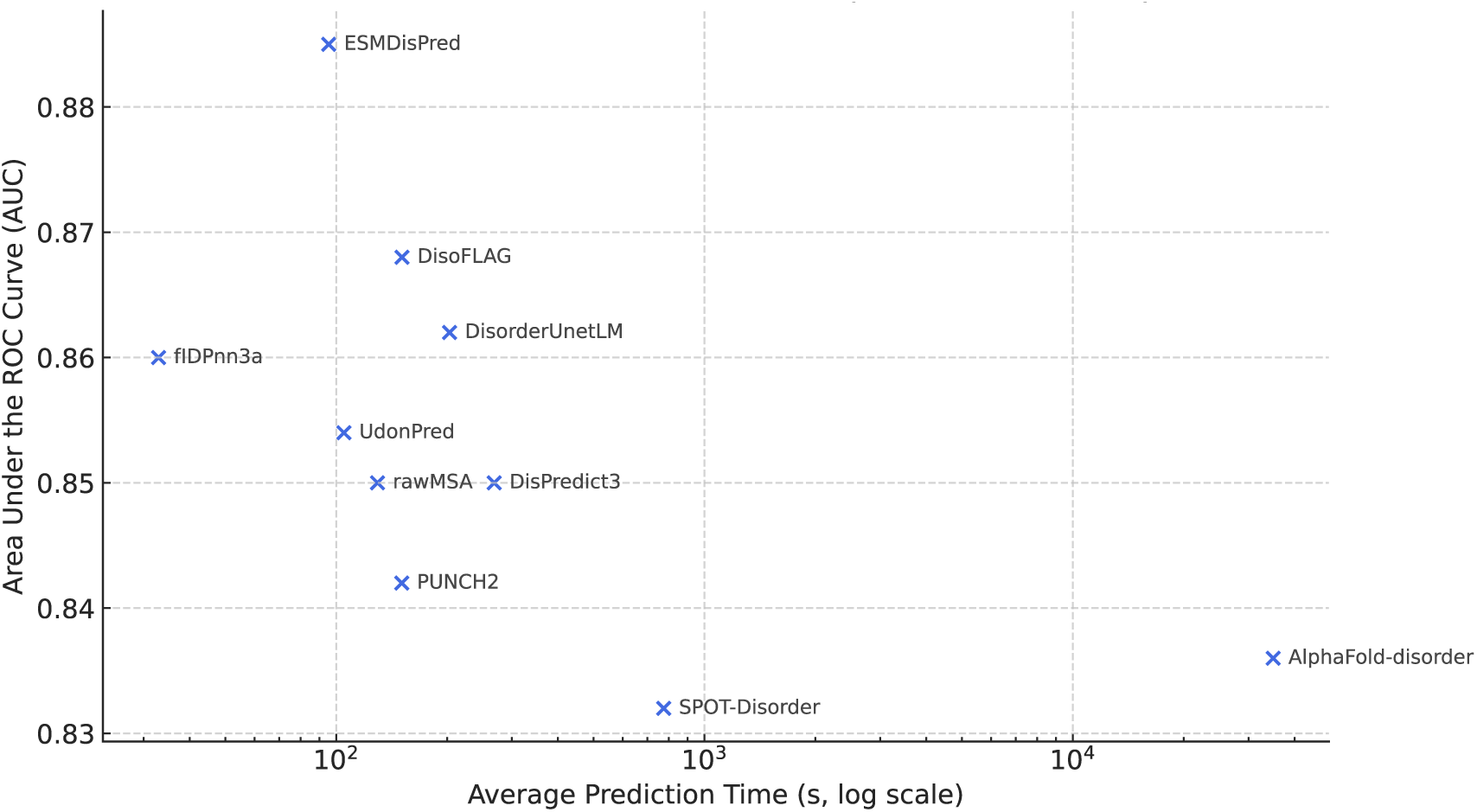
This scatter plot compares the average prediction time (log scale, in seconds) and area under the ROC curve (AUC) for ten leading protein disorder predictors evaluated in the CAID3 challenge. Predictor names are shown next to each point. ESMDisPred achieves the highest AUC with relatively low runtime and shows better performance in both accuracy and efficiency. In contrast, AlphaFold-disorder demonstrates high computational cost (35,000 s) with moderate AUC.

### 1.6 Conclusions

In this work, we introduce ESMDisPred, a structure-aware framework for predicting intrinsically disordered regions in proteins. ESMDisPred integrates fine-tuned embeddings from ESM2 with domain-specific features including terminal annotations, window-based representations, and structural filters. The framework employs a CNN-Transformer hybrid architecture that effectively captures both local sequence motifs and long-range dependencies. This leads to consistent improvements over traditional machine learning algorithms. The empirical evaluation on the Disorder NOX benchmark shows that ESMDisPred achieves substantial performance gains over state-of-the-art methods. These results confirm the framework’s robustness in identifying both fully and partially disordered regions. To further enhance prediction quality, we apply post-processing strategies including Platt scaling, Gaussian smoothing, and hyperparameter optimization. These techniques improve stability in ambiguous cases. Beyond quantitative performance, this study demonstrates the value of combining Transformer-based protein language models with biologically relevant structural context. This integration enhances both predictive accuracy and generalization across diverse protein families.

While ESMDisPred achieves state-of-the-art performance, several directions remain for future investigation. Incorporating AlphaFold-predicted structures could provide lightweight structural cues through per-residue confidence scores (pLDDT) and predicted aligned error (PAE). These scores could help identify flexible segments more accurately. Additionally, modeling the dynamic nature of disorder remains challenging. Residues may transition between ordered and disordered states under different conditions or exhibit low structural confidence. Improving model interpretability through attention visualization and feature attribution will also be critical for uncovering sequence motifs that drive disorder. Despite these remaining challenges, ESMDisPred represents a reliable and practical tool for advancing computational analysis of protein disorders.

## Code availability

The ESMDisPred webserver is available at https://bmll.cs.uno.edu. The code and data related to the development of ESMDisPred can be found here https://github.com/wasicse/ESMDisPred.

## Funding and Acknowledgments

Research reported in this publication was supported by an Institutional Development Award (IDeA) from the National Institute of General Medical Sciences of the National Institutes of Health under grant number P20GM103424-21. Additional support from LA EPSCoR through the NSF EPSCoR SURE Program (NSF Cooperative Agreement No. OIA-2437963) and the Louisiana Optical Network Infrastructure (LONI) for access to high-performance computing resources are gratefully acknowledged.

